# MIRO1 controls energy production and proliferation of vascular smooth muscle cells

**DOI:** 10.1101/2024.08.13.607854

**Authors:** Lan Qian, Olha M. Koval, Benney T. Endoni, Denise Juhr, Colleen S. Stein, Chantal Allamargot, Li-Hsien Lin, Deng-Fu Guo, Kamal Rahmouni, Antentor O. Hinton, E. Dale Abel, Ryan L. Boudreau, Jennifer Streeter, William H. Thiel, Isabella M. Grumbach

## Abstract

**Background:** The outer mitochondrial Rho GTPase 1, MIRO1, mediates mitochondrial motility within cells, but implications for vascular smooth muscle cell (VSMC) physiology and its roles in vascular diseases, such as neointima formation following vascular injury are widely unknown.

**Methods:** Carotid ligation was performed in an in vivo model of selective Miro1 deletion in smooth muscle cells. VSMC proliferation during the cell cycle and molecular mechanisms of smooth muscle cell proliferation were explored in cultured aortic VSMCs by imaging mitochondrial positioning and cristae structure and assessing the effects on ATP production, metabolic function, and interactions with components of the electron transport chain (ETC). MIRO1 expression was also analyzed in human coronary arteries, and its function was assessed via knockdown in human coronary artery VSMCs.

**Results:** MIRO1 was highly expressed in VSMCs within human atherosclerotic plaques. MIRO1 facilitated VSMC proliferation and neointima formation by regulating mitochondrial positioning and PDGF-stimulated ATP production and respiration, critical for cell-cycle progression at G1/S. Deletion of Miro1 disrupted mitochondrial cristae structure, diminished ETC complex I activity, and impaired supercomplex formation. Notably, restoring MIRO1 function with expression of wild type MIRO1 recovered proliferation and ATP production and respiration, whereas a mutant lacking EF hands, which are essential for mitochondrial motility, only partially rescued these effects. MIRO1 knockdown in human coronary artery VSMCs confirmed its pivotal role in mitochondrial function and VSMC proliferation.

**Conclusions:** This study highlights two key mechanisms by which MIRO1 regulates VSMC proliferation. First, it maintains ATP synthesis by preserving mitochondrial cristae integrity. Second, its Ca^2+^-dependent EF hands enable ATP-dependent mitochondrial positioning. By linking mitochondrial motility and energy production to VSMC physiology, these findings position MIRO1 as a critical regulator of vascular remodeling and a potential target for therapeutic interventions.

## INTRODUCTION

Mitochondria play a crucial role in various cellular functions throughout the cardiovascular system [1, 2], yet their specific roles in vascular smooth muscle cells (VSMCs) of systemic arteries remain underexplored, particularly with respect to vasoproliferative diseases such as neointimal hyperplasia. Insights into these processes have largely stemmed from extending findings on cytosolic pathways. For instance, neointima formation has been shown to be inhibited either by promoting apoptosis via mitochondrial pathways [3–5] or by reducing mitochondrial ROS production through overexpression of uncoupling protein 2 or superoxide dismutase 2 [6–8].

Mitochondria dynamically adapt to cellular changes through fission and fusion, which modulate VSMC proliferation, migration, and neointima formation [9–11]. Moreover, over a decade ago Chalmers et al. showed that interfering with mitochondrial movement blocks VSMC proliferation [12]. Despite this, the molecular regulators governing mitochondrial motility in VSMCs have remained understudied.

The outer mitochondrial membrane GTPase MIRO1 is essential for orchestrating mitochondrial positioning in many cell types, including neurons, lymphocytes, and various cancer cell lines [13–16]. MIRO1 contains two canonical EF-hand domains that are flanked by two GTPase domains and a C-terminal transmembrane domain that anchors MIRO1 to the outer mitochondrial membrane [17]. MIRO1 associates with trafficking kinesin-binding proteins (TRAKs), linking mitochondria to microtubules and myosin XIX [13–15, 18]. These interactions are regulated by changes in intracellular Ca^2+^ levels: when MIRO1 is not bound by Ca^2+^/calmodulin, it links mitochondria to microtubules; when the EF hands are bound by Ca^2+^, a conformational change causes mitochondria to dissociate from microtubules, arresting their movement [13]. In addition to playing a role in promoting mitochondrial motility, MIRO1 is believed to control mitophagy through phosphorylation by the Pink/Parkin complex [19, 20], and to facilitate the formation of mitochondrial cristae via associations with the mitochondrial contact site (MICOS) complex and the mitochondrial intermembrane space bridging (MIB) complex [21].

Reports of MIRO1’s diverse roles have prompted some investigations into its role in human diseases. For example, Miro1 mutations identified in humans (for example p.R272Q) have been linked genetically and pathophysiologically to Parkinson’s disease [19, 20, 22]. Nevertheless, few studies have investigated its role in cardiovascular disease. Cardiomyocytes isolated from neonatal rats lacking Miro1 were protected against phenylephrine-induced cardiomyocyte hypertrophy and mitochondrial fission [23]. However, its specific contributions to vascular pathologies were less well understood, with data from our laboratory highlighting an important role for MIRO1 in VSMC migration [24]. Emerging evidence positions MIRO1 as a regulator of cell proliferation in fibroblasts [16, 25, 26]. Our group recently reported that cyclic changes in formation of mitochondrial ER contact sides (MERCS) that enable the transfer of Ca^2+^ and other metabolites to mitochondria are required for cell proliferation and controlled by MIRO1 [25].

The goal of this study was to determine whether MIRO1 controls VSMC proliferation and, if so, to uncover the underlying mechanisms involved. We investigated the effects of MIRO1 manipulation on VSMC proliferation and the potential mechanism by which MIRO1 affects cell cycle progression, mitochondrial motility, ATP production, and respiration. Additionally, we used a transgenic mouse model in which Miro1 was deleted specifically in VSMCs to test its effects on neointima formation in vivo.

## RESULTS

### Loss of MIRO1 blocks neointima formation

To study the effects of MIRO1 on smooth muscle cell biology after vascular injury, we generated a transgenic model in which MIRO1 is selectively deleted in smooth muscle cells after administration of tamoxifen (SM-MIRO1^-/-^). Reduced levels of MIRO1 mRNA and protein in carotid arteries and the aorta were confirmed by immunohistochemistry, western blotting, and quantitative real time PCR (**Figure 1A**, **Figure 1- Figure supplement 1A, B**). Mice of both the wild type and SM-MIRO1^-/-^ genotypes were fed a high-fat diet for three weeks, and then vascular injury was induced by ligation of the common carotid artery. Three weeks after ligation, the mice were euthanized and neointima size was assessed. Neointima formation was robust in wild type mice and significantly reduced in SM-MIRO1^-/-^ mice (**Figure 1B**). The neointimal area was smaller in the arteries from the SM-MIRO1^-/-^ mice than in those of their wild type counterparts.

**Figure 1.**
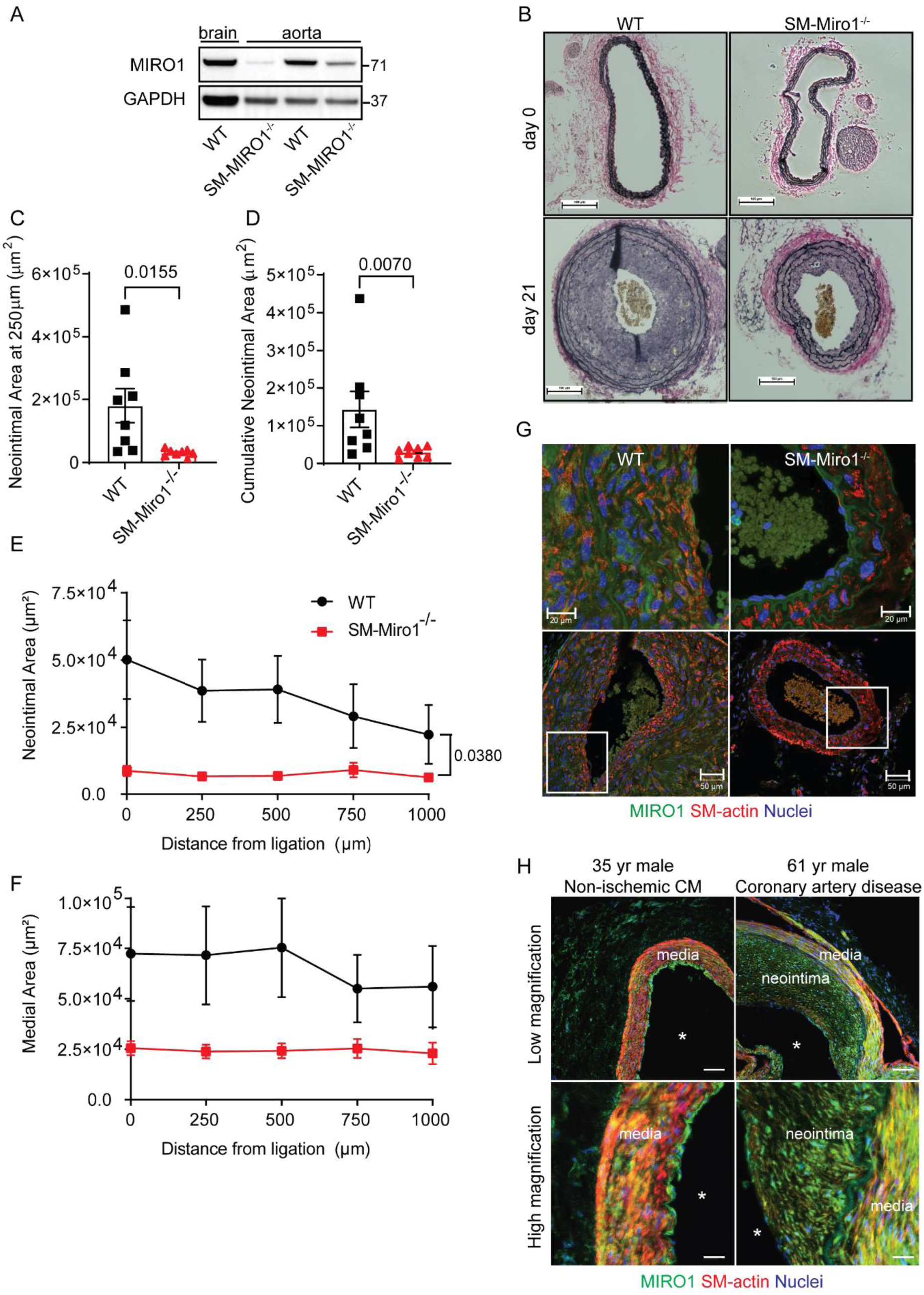
Loss of Miro1 blocks neointima formation. (A) Immunoblots for MIRO1 in lysates from brain and aorta isolated from WT and SM-Miro1^-/-^mice. (B) Verhoeff-Van Gieson staining in the unligated and ligated common carotid arteries of wild type and SM-Miro1^-/-^ mice, at 21 days post ligation. Scale bar = 100 µm. (C) Neointimal area in ligated carotid arteries at 250 µm from the bifurcation. Neointimal area was determined by subtracting the luminal area from the area defined by the internal elastic lamina. (D) Cumulative neointimal area, calculated from all neointimal areas within 1000 µm of the bifurcation. (E) Neointimal area at the indicated distances from the site of ligation. (F) Medial area at the indicated distances from the site of ligation. The medial area was determined by subtracting the area defined by the internal elastic lamina from the area defined by the external elastic lamina. (G) Immunofluorescence of MIRO1 expression in the mouse carotid artery. MIRO1, green; SM-actin, red; Nuclei, DAPI, blue. Scale bar: 20 µm in upper images; 50 µm in lower images. Upper images are magnifications of the areas labeled with a box in the lower images. (H) Immunofluorescence of MIRO1 in the left anterior descending artery of a healthy subject and that of a patient with coronary artery disease. MIRO1, green; SM-actin, red; Nuclei, DAPI, blue. Scale bar: 80 µm in upper, 20 µm in lower images. Statistical analyses were performed using unpaired t-test (C), Mann-Whitney test (D) and two-way ANOVA (E, F).

Moreover, the neointimal area was decreased at 250 μm from the ligation site and the cumulative neointimal area over the first one mm of the carotid artery proximal to the ligation (**Figure 1C-F**). Immunohistochemical analysis in mouse carotid arteries after ligation revealed expression of MIRO1 in VSMCs of the neointima (**Figure 1G**). Lastly, we confirmed that MIRO1 is present in human coronary arteries. In a subject without a medical history of coronary disease, MIRO1 was present in the media. In human subjects with atherosclerotic disease, MIRO1 was expressed in both the media and the neointima, in cells also positive for smooth-muscle actin (**Figure 1H, Figure 1- Figure supplement 2**). Moreover, in the atherosclerotic plaque, we detected the colocalization of MIRO1 with the proliferation marker Ki67. In some cells, both were colocalized with smooth-muscle actin (**Figure 1- Figure supplement 3**).

### Loss of MIRO1 reduces the proliferation of smooth muscle cells

To determine the relevance of MIRO1 to cell proliferation, VSMCs were explanted from the aortas of SM-MIRO1^-/-^ mice and cultured. Platelet-derived growth factor (PDGF) was used to enhance cell proliferation. Cell number was assessed 3 days after plating. In the case of wild type cells, the number was approximately 80% greater for those treated with PDGF than for those not treated with PDGF. In the case of SM-MIRO1^-/-^ VSMCs, cell counts were lower in non-treated cells and in cells following PDGF-treatment (**Figure 2- Figure supplement 1A**).

Given the poor proliferation of VSMCs from SM-MIRO1^-/-^ mice, in later experiments we used VSMCs isolated from MIRO1^fl/fl^ mice and infected them with adenovirus expressing cre recombinase (Ad Cre; these cells are henceforth denoted as Miro1^-/-^). As negative controls, we used cells isolated from the same mice infected with an empty vector control adenovirus (Ad EV). In this model, cre-mediated MIRO1 recombination was efficient (**Figure 2A, Figure 2-Figure supplement 1B-D**). At 72 h after plating in growth medium, the number of Miro1^-/-^VSMCs was significantly lower than that of wild type cells, in growth media and following PDGF treatment (**Figure 2B**). Cells were synchronized by serum starvation for 48 h and FACS analysis was performed to identify the phase of the cell cycle during which the growth delay caused by a lack of MIRO1 occurred (**Figure 2C-F**). At baseline (0 h), no differences in the cell cycle distribution were observed across the experimental groups (**Figure 2D**). At both 24 h and 48 h, significant differences in percentage of cells in G1 and S phase were present. At 24 h, there were significantly fewer wild type than MIRO1^-/-^ VSMCs in G1 phase and more wild type than MIRO1^-/-^ VSMCs in S phase. However, at 48 h, the percentage of wild type cells in G1 was more than 80%, similar to the percentage at 0 h, suggesting that the cell cycle was completed. In contrast, at 48 h, more MIRO1^-/-^ VSMCs remained in S phase and fewer were in G1 phase than at 0 h, indicating a delay in cell cycle (**Figure 2D-F**).

**Figure 2.**
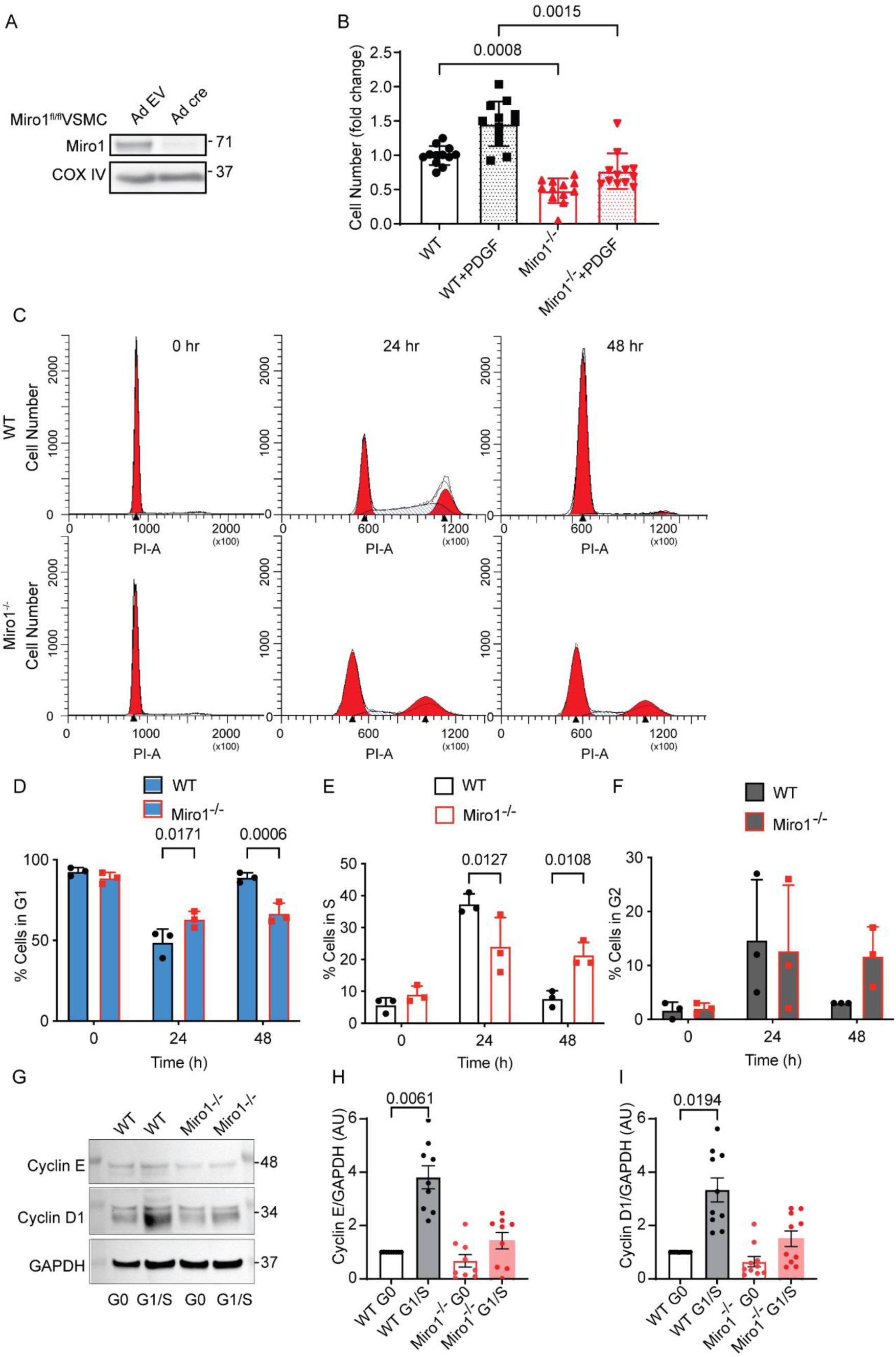
Loss of Miro1 reduces G1/S transition and VSMC proliferation. (**A**) MIRO1 levels in mitochondrial fractions of VSMCs from Miro1^fl/fl^ mice transduced with adenovirus expressing Cre recombinase or control adenovirus, as determined by immunoblotting. (**B**) Number of VSMCs from Miro1^fl/fl^ mice following transduction with adenovirus expressing cre recombinase (MIRO1^-/-^) or control (WT) adenovirus, at 72 h after incubation with PDGF (20 ng/ml) or control (saline). (**C**) DNA content of synchronized WT and MIRO1^-/-^ VSMCs, as assessed by fluorescence-activated cell sorting (FACS). Times are 0, 24, and 48 h after release from growth arrest for 48h in FBS-free media, and then at 24 and 48 h after release from arrest with media containing 10% FBS. Analysis compared differences between genotypes. (**D-F**) Quantification of cell-cycle phase distribution of WT and MIRO1^-/-^ VSMCs shown in C. (**G**) Immunoblots for cyclin E and D1 as markers of the indicated cell cycle phases in whole-cell lysates of WT and Miro1^-/-^ VSMCs following synchronization by serum starvation, at G0 (after growth arrest for 48 h in FBS-free media), and G1/S (after release from growth arrest with media containing 10% FBS for 24 h). (**H, I**) Quantification of Cyclin D1 and E levels on immunoblots like those shown in panel G. Statistical analyses were performed using Kruskal-Wallis test (B) and two-way ANOVA (D-F), and Friedman test (H, I).

Immunoblotting for cyclin D1 and cyclin E in VSMCs incubated in serum-containing growth media for 24 h revealed lower protein levels in the G1/S transition in MIRO1^-/-^ VSMCs (**Figure 2G-I**), consistent with cell cycle delay and similar to the results of the FACS analysis.

### MIRO1’s EF hands are required for mitochondrial motility and cell proliferation

We also evaluated the extent to which the reconstitution of MIRO1 in MIRO1^-/-^ VSMCs rescued cell proliferation. Expressing wild type MIRO1 in MIRO1^-/-^ VSMCs at levels similar to those in wild type cells normalized proliferation, whereas expressing a MIRO1 mutant lacking the EF hands at same levels only partially rescued cell proliferation (**Figure 3A-C**). These data demonstrate that MIRO1 is required for cell proliferation, that it affects early cell cycle progression during the G1/S-phase, and that the EF hands of MIRO1 are necessary for normal cell proliferation. The EF hands of MIRO1 mediate the attachment to microtubules in neurons and without them, mitochondria do not move normally within axons [13].

**Figure 3.**
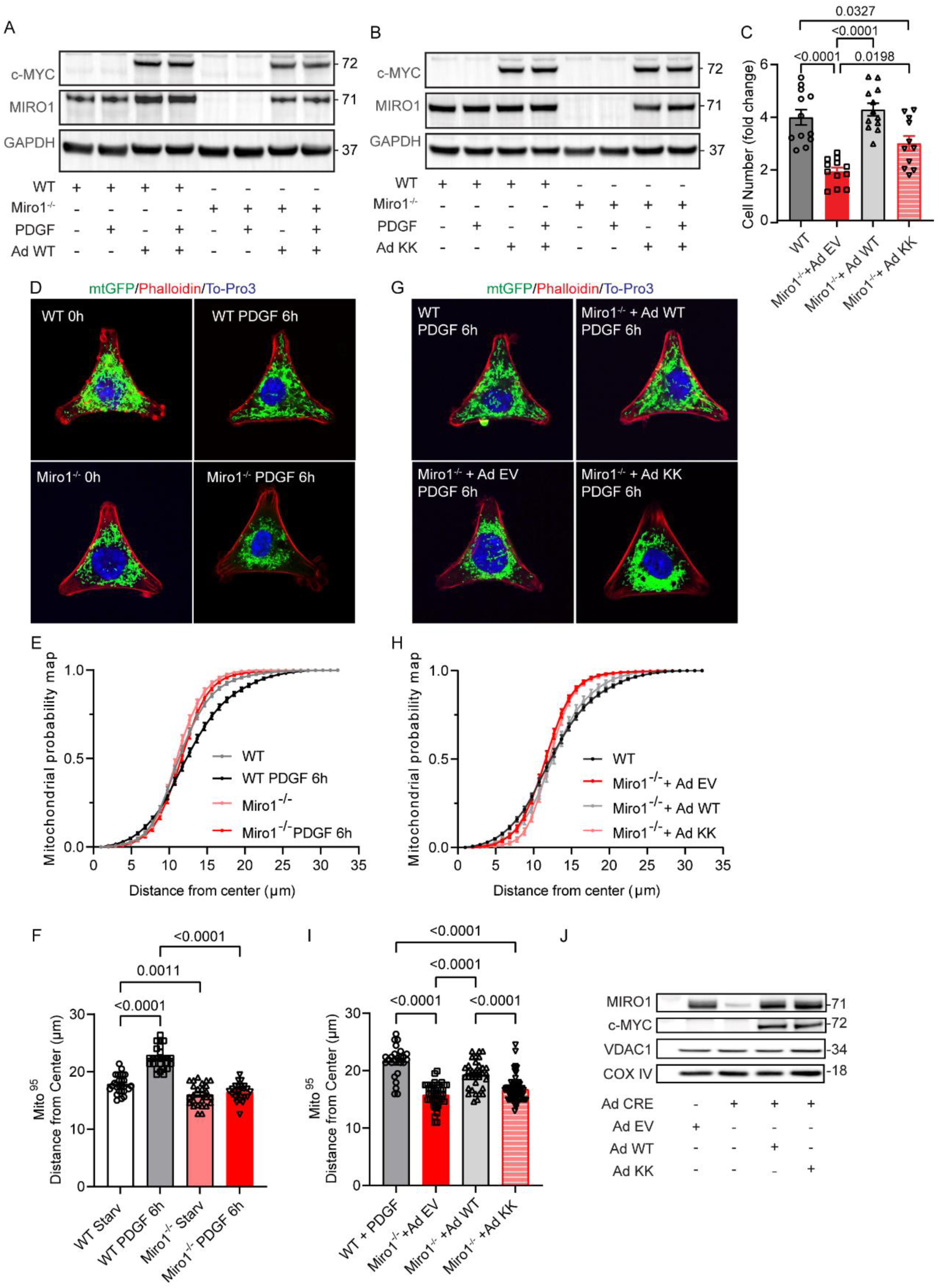
Loss of EF hands in MIRO1 reduces mitochondrial motility and cell proliferation. (**A, B**) Immunoblots for MIRO1 and c-MYC in WT and Miro1^-/-^ VSMCs transduced with adenovirus expressing MIRO1-WT (A) and MIRO1-KK (B). (C) Number of WT and Miro1 ^-/-^ VSMCs transduced with adenovirus expressing MIRO1-WT (Ad WT), MIRO1-KK (Ad KK), or control adenovirus (empty vector, Ad EV) and treated with PDGF (20 ng/ml) for 72 h. (D) Representative confocal images of WT and Miro1^-/-^ VSMCs grown on Y-shaped adhesive micropatterns (CYTOOchips^TM^), with VSMCs synchronized by serum starvation for 24 h (0 h timepoint), followed by change to medium containing 10% FCS and PDGF (20 ng/ml) for 6h (Phalloidin, red; mitochondrial GFP, green; To-Pro-3, blue). (E) Mitochondrial probability map. The cumulative distribution of mitochondria was assessed for images as in (D) by modified Sholl’s analysis. Data are plotted by growth conditions (as in A). (F) Mito^95^ values, defined as the distance from the center of the CYTOOchips^TM^ at which 95% of the mitochondrial signal is found, under the conditions used in (A). (G) Representative confocal images of WT and Miro1^-/-^ VSMCs grown on CYTOOchips^TM^. Miro1^-/-^ VSMCs were transduced with adenovirus expressing MIRO1-WT or MIRO1-KK for 72 h before being seeded onto CYTOOchips^TM^. The cell cycle was synchronized by serum starvation for 24 h (not shown); the cells were subsequently treated with medium containing 10% FCS and PDGF (20 ng/ml) for 6 h. (H) Mitochondrial probability map. The cumulative distribution of mitochondria was assessed for images shown in (G) by modified Sholl’s analysis. Data are plotted by growth conditions (as in B). (I) Mito^95^ values, as defined in (F) but under the conditions used in (B). (J) Immunoblots for MIRO1 and c-Myc in WT and Miro1^-/-^ VSMCs transduced with adenoviruses expressing MIRO1-WT, MIRO1-KK, or with control adenovirus. Statistical analyses were performed by one-way ANOVA (C, F) and Kruskal-Wallis test (I).

To determine whether MIRO1^-/-^ cells can form fused mitochondrial networks sufficient to support the G1/S-phase, we assessed mitochondrial morphology at different points of the cell cycle. Analysis of mitochondrial form factor, defined as the ratio of mitochondrial length to width, revealed morphological changes in wild type cells, characterized by an increase in fusion at 24 h and 48 h (**Figure 3- Figure supplement 1A**). In contrast, MIRO1^-/-^ cells exhibited no significant alterations in mitochondrial morphology. No significant reduction in mitochondrial DNA copy number was detected MIRO1^-/-^ VSMCs compared to wild type cells (**Figure 3- Figure supplement 1B**).

To determine the extent to which MIRO1 contributes to mitochondrial motility in VSMCs, we plated wild type or MIRO1^-/-^ VSMCs on microchips (CYTOOchip^TM^) with Y-patterns. In wild type VSMCs, after cell cycle arrest caused by serum starvation, mitochondria were concentrated around the nucleus. At 6 h after treatment in growth media supplemented with PDGF, the mitochondria were distributed throughout the cell, including near the cell edges (**Figure 3D-F**). In the MIRO1^-/-^ VSMCs, however, the mitochondria remained near the nucleus. Additionally, by live-cell fluorescence confocal microscopy, wild type cells exhibited dynamic reorganization of the mitochondrial network, whereas MIRO1^-/-^ VSMCs displayed minimal mitochondrial movement, characterized only by limited oscillatory behavior without network remodeling (**Figure 3- Video supplement 1**). Reconstitution of MIRO1^-/-^ VSMCs with wild type MIRO1 fully rescued the mitochondrial distribution at 6 h after PDGF treatment (**Figure 3G-J**). In contrast, after transduction of MIRO1^-/-^ VSMCs with the MIRO1 mutant KK (MIRO1-KK, Ad KK), which lacks functional EF hands, the mitochondria were irregularly and unequally distributed. These data demonstrate that MIRO1 is required for both cell proliferation and coordinated mitochondrial motility within the cell. In the absence of EF hands, proliferation and motility is partially restored, suggesting that MIRO1 supports proliferation through a mechanism beyond control of motility.

We sought to further elucidate the relationship between ATP production, mitochondrial motility, and VSMC proliferation. First, we determined whether mitochondrial ATP production is necessary for mitochondrial motility or VSMC proliferation. For this purpose, we added oligomycin, an inhibitor of ATP synthesis, to proliferating wild type VSMCs for 72 h. This treatment reduced the number of VSMCs (**Figure 4A**) as well as in the intracellular ATP level (**Figure 4B**). To evaluate the effect of ATP production by mitochondria on their motility, we assessed the mitochondrial distribution in wild type VSMCs plated on micropatterned CYTOOchips. Mitochondrial motility was reduced when ATP production was inhibited by oligomycin (**Figure 4C-F**). To investigate whether blocking mitochondrial motility affects intracellular ATP levels, we performed control experiments with Nocodazole, which inhibits microtubule assembly. This treatment effectively arrested mitochondrial motility, but ATP levels were not affected (**Figure 4G, H).** These findings support that mitochondrial respiration is required for VSMC proliferation as well as mitochondrial motility along microtubules and that MIRO1 controls both. In contrast, mitochondrial motility is not required for optimal cellular ATP production.

**Figure 4.**
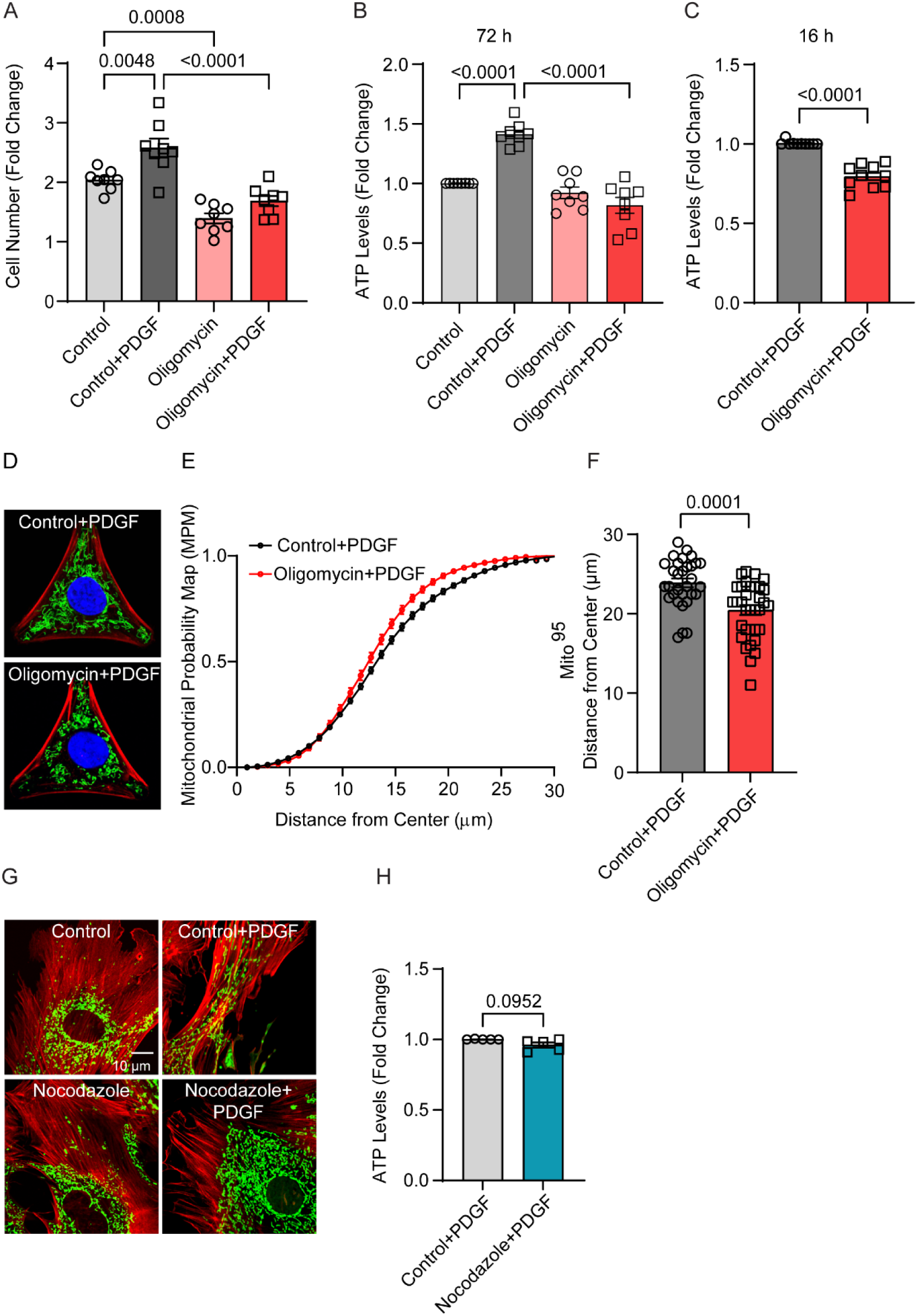
Inhibition of mitochondrial ATP production reduces VSMC proliferation, but inhibiting mitochondrial motility does not affect ATP levels. (**A**) Number of WT VSMCs after 72-h treatment with either PDGF (20 ng/ml) and/or oligomycin (1 µM). (**B, C**) ATP levels in WT VSMCs after treatment with PDGF (20 ng/ml) or with PDGF in addition to oligomycin (1 µM) for 72 h (B) or 16 h (C). (D) Representative confocal images of WT VSMCs on CYTOOchips^TM^ with Y-shaped micropatterns, following 16-h treatment with PDGF (20 ng/ml) and oligomycin (1 µM). Mitochondria, green; phalloidin, red; DAPI, blue. (E) Mitochondrial probability map. The cumulative distribution of mitochondria was assessed for images as in (D) by modified Sholl’s analysis. (F) Mito^95^ values, defined as the distance from the center of the Y-shaped pattern at which 95% of the mitochondrial signal is found for growing cells like in (D). (G) Representative confocal images of WT VSMCs infected with mitoGFP and stained with phalloidin. Cells were synchronized by serum starvation for 48 h and then released from starvation by replacing the medium with one containing 10% FBS and PDGF (20 ng/ml). Some samples were treated with nocodazole (1 µM). Images were taken immediately after growth media (Control) or growth media with nocodazole was added (Nocodazole) and after incubation for 16 h (Control + PDGF, Nocodazole + PDGF). (H) ATP levels in WT VSMCs after 16-h treatment in growth media with PDGF or growth media supplemented with PDGF and Nocodazole (1 µM). Statistical analyses were performed by one-way ANOVA (A, B) or Mann-Whitney test (C, F, H).

### Loss of MIRO1 leads to impaired metabolic activity and reduced proliferative capacity

Our findings that mitochondrial motility is controlled by MIRO1 and depends on mitochondrial ATP production imply that Miro1 deletion causes a defect in metabolic activity. Given that cell proliferation is impaired at the early stages of the cell cycle in VSMCs with MIRO1 deletion and that intracellular ATP demands are high during the G1/S transition [27, 28], we measured intracellular ATP levels in cells after 16 h of PDGF treatment. MIRO1^-/-^ samples exhibited reduced ATP levels accompanied by elevated concentrations of ADP and AMP (**Figure 5A-C**). As a result, both ATP/ADP and ATP/AMP ratios were significantly lower in MIRO1^-/-^ cells compared to wild type, indicating impaired cellular energy homeostasis (**Figure 5B, C)**.

**Figure 5.**
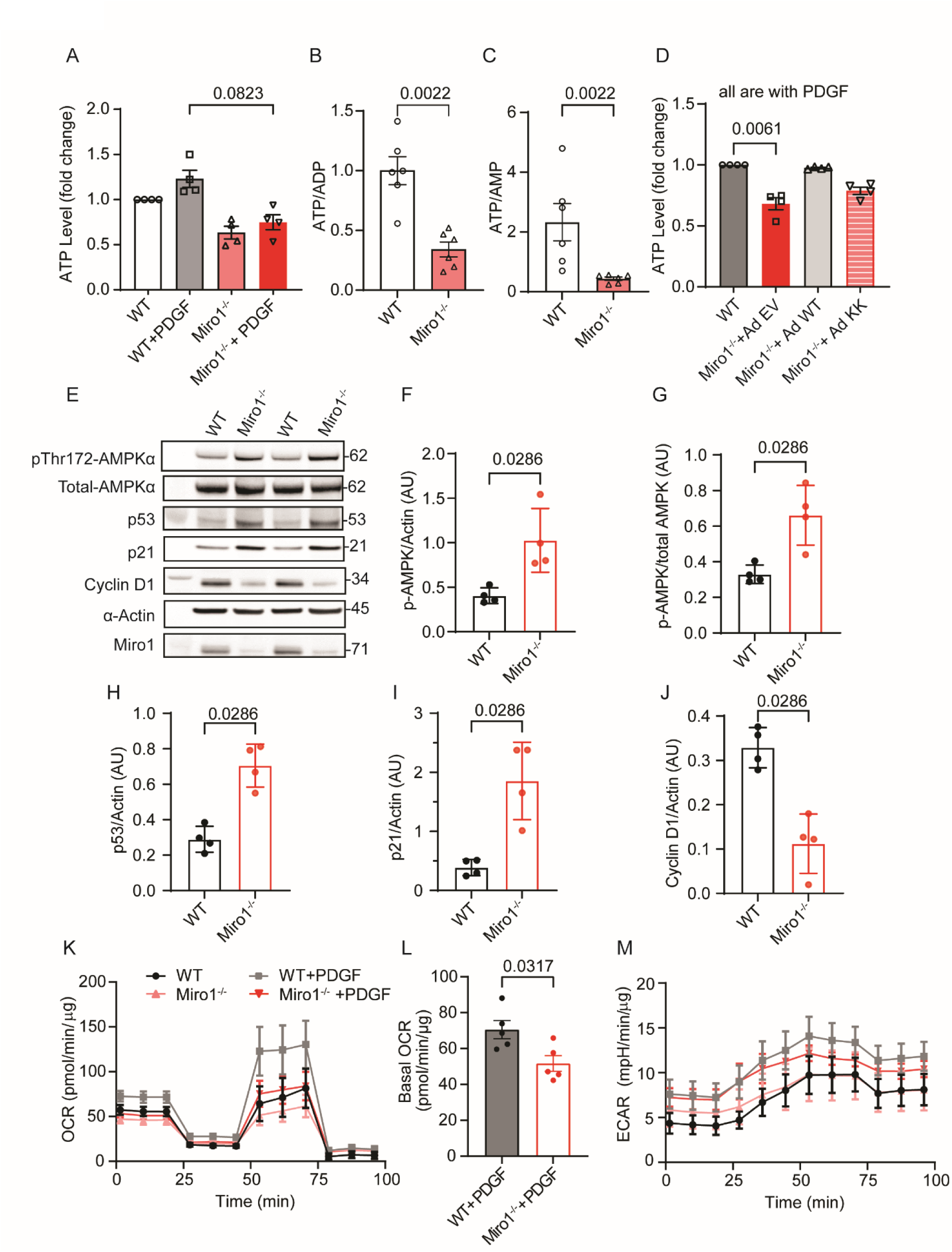
Loss of Miro1 impairs mitochondrial metabolic activity and is associated with decreased proliferative capacity. (A) Intracellular ATP levels in WT and Miro1^-/-^ VSMCs after treatment with PDGF (20 ng/ml for 16 h) or control (no PDGF), normalized to levels in control WT VSMCs (no PDGF). (**B, C**) ATP/ADP (B) and ATP/AMP (C) ratios assessed in WT and Miro1^-/-^ VSMCs.(**D**) Intracellular ATP levels in WT and Miro1^-/-^ VSMCs transduced with adenovirus expressing MIRO1-WT (Ad WT) or MIRO1-KK (Ad KK) or control adenovirus (empty vector, Ad-EV) after treatment with PDGF (20 ng/ml for 16 h) or control (no PDGF), normalized to ATP levels in control WT VSMCs (no PDGF). (**E**) Immunoblot of lysates from WT and Miro1^-/-^ VSMCs for markers of the metabolic state (phosphorylated (p-Thr172) and total AMPKα), and cell cycle progression (p53, p21 and cyclin D1) at 48 h after incubation in serum-free medium (cell-cycle arrest). (**F-J**). Quantification of immunoblot signal in samples shown in (E) for (F) phosphorylated (p-Thr172) AMPKα normalized to actin, (G) phosphorylated (p-Thr172) AMPKα normalized to AMPKα, (H) p53, (I) p21, and (J) cyclin D1, all normalized to actin. (K) Oxygen consumption rate (OCR), as determined by Seahorse, for WT and Miro1^-/-^ VSMCs with and without PDGF treatment (20 ng/ml for 16 h). (L) Quantification of basal OCR for WT and Miro1^-/-^ VSMCs treated with PDGF (n=5). (M) Extracellular acidification rate (ECAR), as determined by Seahorse, for WT and Miro1^-/-^VSMCs with and without PDGF treatment (20 ng/ml for 16 h). Statistical analyses were performed by Friedman test (A, D), and Mann-Whitney (B, C, F-J, L).

Additionally, transduction with MIRO1-WT normalized ATP production, whereas transduction with MIRO1-KK did not (**Figure 5D**). AMP kinase is an intracellular indicator of the metabolic state and regulates the expression of cyclins D and E. Thus, we determined AMP kinase activation by immunoblotting. In MIRO1^-/-^ VSMCs, the phosphorylation of AMP kinase was elevated, indicating intracellular energy deficiency (**Figure 5E-G**). AMP kinase suppresses cell cycle progression through phosphorylation and stabilization of p53 and subsequent induction of the p21, which leads to cell cycle arrest in G1 phase. In MIRO1^-/-^ VSMCs, the p53 and p21 levels were increased and cyclin D1 decreased (**Figure 5H-J**), supporting that MIRO1 is required for energy production and cell-cycle progression in the G1/S phase. We also measured oxygen consumption rates in wild type and Miro1^-/-^ VSMCs by Seahorse respirometry. We detected decreased basal respiration in MIRO1^-/-^ VSMCs treated with PDGF compared to wild type (**Figure 5K, L**). However, the rate of extracellular acidification, a measure of glycolysis, was not significantly affected in MIRO1^-/-^ VSMCs (**Figure 5M**), indicating a lack of compensatory upregulation of glycolysis. These findings were further confirmed by Seahorse metabolic assays on wild type, MIRO1^-/-^, and MIRO1^-/-^ VSMCs expressing MIRO1 wild type (by adenoviral transduction with Ad WT), MIRO1KK (Ad KK), and a MIRO1 mutant lacking the C-terminal transmembrane domain (Ad ΔTM, **Figure 5- Figure supplement 1A-D**). Only the expression of MIRO1 wild type in MIRO1^-/-^ VSMCs restored mitochondrial respiration. Neither mitochondrial respiration nor glycolytic rate were affected by Nocodazole treatment (**Figure 5- Figure supplement 1B, D**).

### MIRO1 controls electron transport chain (ETC) activity

We sought to establish the mechanism by which MIRO1 controls ATP production. Previous findings in embryonic fibroblasts have suggested that MIRO1 associates with both the mitochondrial contact site (MICOS) complex and the mitochondrial intermembrane space bridging (MIB) complex, which collectively control the folding of mitochondrial cristae [21]. Moreover, we recently reported that MIRO1 regulates the formation of mitochondria-ER contact sites (MERCS) during the cell cycle and facilitates the Ca^2+^ transfer to mitochondria [25]. We assessed the effect of MIRO1 deletion on the formation of mitochondrial cristae in VSMCs by transmission electron microscopy. Consistent with findings in fibroblasts [21], we found that in Miro1^-/-^ VSMCs, the cristae were less dense than those in wild type controls, and their morphology was distorted (**Figure 6A-C**). Next, we established the associations of wild type MIRO1, MIRO1-KK, and a MIRO1 mutant lacking the C-terminal transmembrane domain (MIRO1-ΔTM) with components of the electron transport chain and the MIB/MICOS complex.

**Figure 6.**
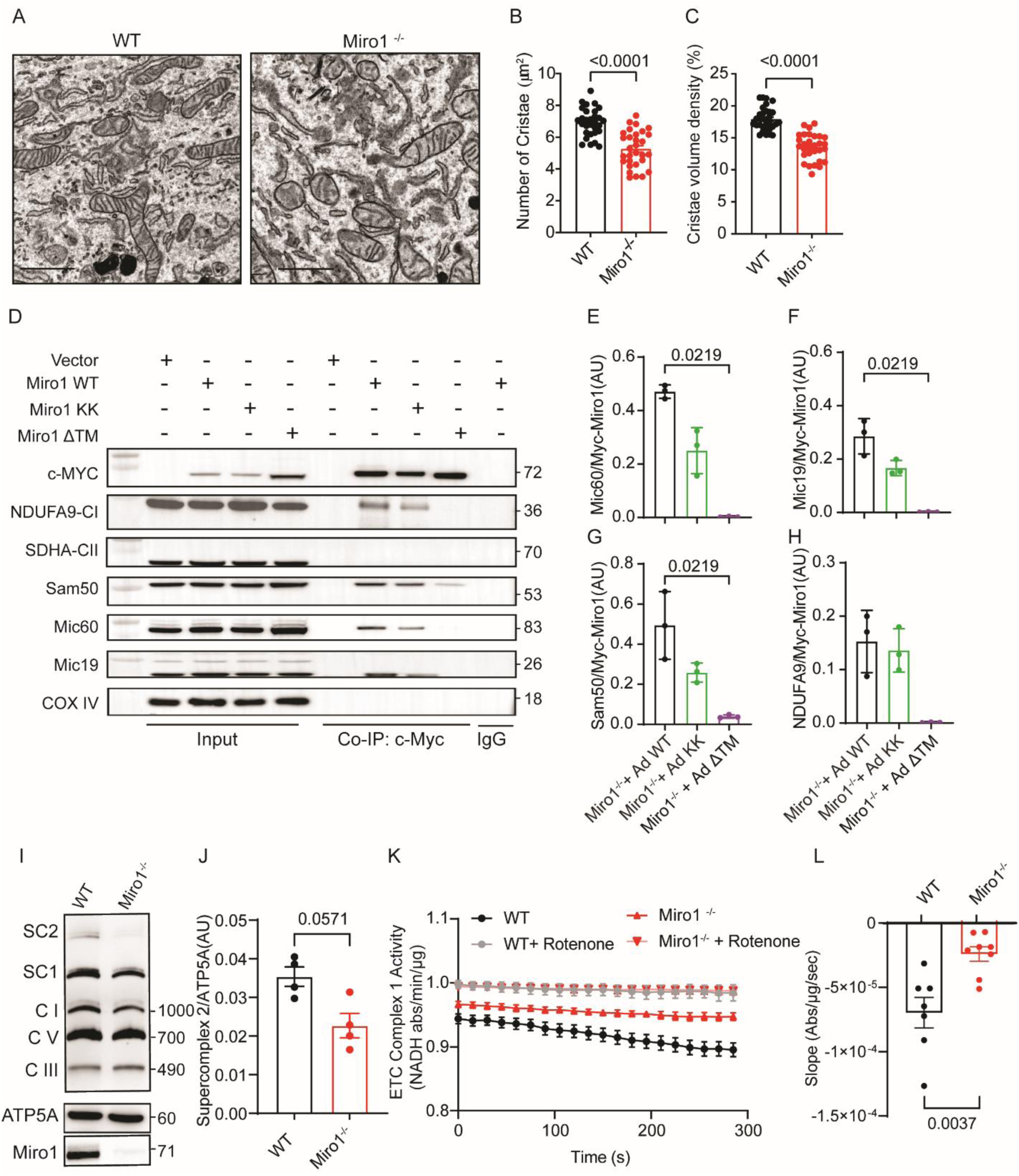
Loss of Miro1 leads to reduced ETC activity under growth conditions. (A) Transmission electron microscopy images of WT and Miro1^-/-^ VSMCs. (**B, C**) Quantification of (B) the number of mitochondrial cristae and (C) the volume density of the mitochondrial cristae in WT and Miro1^-/-^ VSMCs. (**D**) Levels of c-Myc tagged MIRO1-WT (Ad WT), c-Myc tagged MIRO1-KK (Ad KK), or MIRO1-ΔTM (Ad ΔTM) and proteins of the MIB/MICOS complex in HEK cells following pull-down assay, as determined by immunoblotting. C-Myc tagged Miro1 constructs were expressed in HEK cells for 72 h before cell lysis and pull-down were performed. (**E-H**) Quantification of (E) MIC60, (F) MIC19, (G) SAM 50, and (H) NDUFA9 from (D), adjusted for immunoprecipitated c-Myc-tagged MIRO1 as in (D). (**I**) Levels of mitochondrial supercomplex and ETC subunits in WT and Miro1^-/-^ VSMCs, as determined by blue-native poly-acrylamide gel electrophoresis (BN-PAGE). (**J**) Quantification of supercomplex 2 in (I). (**K**) Quantification of activity of ETC complex 1 in WT and Miro1^-/-^ VSMCs, as determined by the decrease in the rate of absorbance at 340 nm with and without rotenone incubation for 10 min. (**L**) Quantification of activity of ETC complex 1, plotted as the difference between absorbance curve slopes with and without rotenone (as in K). Statistical analyses were performed by unpaired t-test (B, C), Kruskal-Wallis (E-H), Mann-Whitney (J, L).

Pull-down assays in HEK293T cells revealed that MIRO1 interacts with multiple components of the mitochondrial intermembrane space bridging (MIB) complex, including the SAM complex subunit Sam50 and the MICOS complex subunits Mic60 and Mic19 (Figure 6D–G), as well as with the NDUFA9 subunit of electron transport chain complex I (Figure 6H). The transmembrane domain by which MIRO1 is attached to mitochondria was necessary for the association with all proteins, and in the absence of the EF hands the association with all proteins was weaker. The expression of proteins of the MIB/MICOS complex was not affected by MIRO1 deletion (**Figure 6- Figure supplement 1A**).

Immunoblotting for subunits of all five electron transport chain complexes did not reveal a significant difference in protein levels between wild type and MIRO1^-/-^ VSMCs (**Figure 6- Figure supplement 1B-G**). An in-gel Complex V activity assay was performed to evaluate the enzymatic activity of mitochondrial ATP synthase within a native gel following electrophoresis. To normalize the activity signal, a Blue Native PAGE of the same samples was immunoblotted for the ATP5F1 subunit. A modest yet statistically significant reduction in Complex V activity was observed in MIRO1^-/-^ samples (**Figure 6- Figure supplement 1H-K**).

Because the formation of mitochondrial cristae affects the formation of the MIB/MICOS supercomplex and the activity of the ETC chain, we assessed the abundance of mitochondrial supercomplexes in blue native gels (**Figure 6I, J**). Supercomplex 2 was decreased in MIRO1^-/-^VSMCs relative to wild type VSMCs (**Figure 6J**). We also tested the activity of ETC complex I by measuring the consumption of NADH. In MIRO1^-/-^ cells, the activity of this complex was significantly decreased (**Figure 6K, L**). Overexpression of wild type MIRO1 fully restored enzymatic activity, whereas MIRO1-KK provided partial rescue (**Figure 6- Figure supplement 2A-C, E**). In contrast, the mutant MIRO1-ΔTM failed to restore activity and resembled the MIRO1 knockout phenotype (**Figure 6- Figure supplement 2D, E**). ETC activity was abolished by incubation with Rotenone (**Figure 6- Figure supplement 2F).** These data demonstrate that MIRO1 associates with MIB/MICOS. Moreover, MIRO1 promotes the formation of mitochondrial supercomplexes and the activity of ETC complex I.

### MIRO1 knockdown in human coronary smooth muscle cells impairs proliferation, mitochondrial motility, and ETC activity

To determine if these findings also apply to human cells, MIRO1 was knocked down in human coronary smooth muscle cells (**Figure 7A, B**). Cell counts 72 hours after plating revealed a significant decrease in the number of cells with MIRO1 knockdown, particularly in those treated with PDGF (**Figure 7C**). Additionally, in VSMCs plated on microchips (CYTOOchipTM) with Y-patterns and treated with PDGF, MIRO1 knockdown led to a concentration of mitochondria near the nucleus, similar to mouse MIRO1^-/-^ VSMCs. In control conditions, the mitochondria were distributed throughout the cell, including near the edges (**Figure 7D-F**). Intracellular ATP levels were reduced with MIRO1 knockdown (**Figure 7G-I**). Finally, the activity of ETC complex I, measured by NADH consumption, was significantly decreased in MIRO1^-/-^ cells (**Figure 7J, K**). These results indicate that the effects of MIRO1 on proliferation and metabolism in VSMCs are conserved among species and could be clinically relevant for human disease.

**Figure 7.**
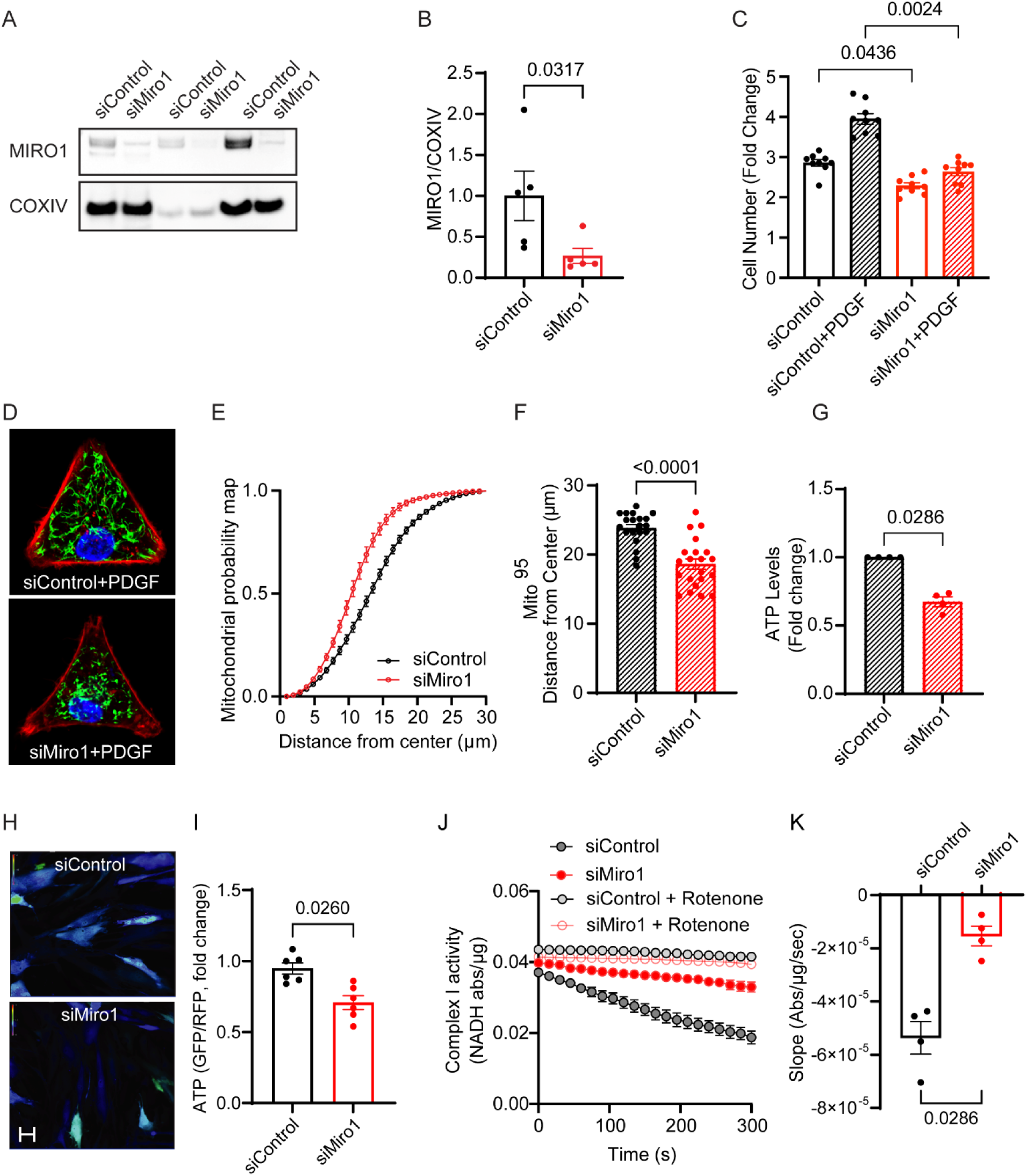
MIRO1 knockdown in human coronary artery smooth muscle cells inhibits mitochondrial motility, mitochondrial bioenergetics, and proliferation. (A) Immunoblot for MIRO1 and COX IV in mitochondrial fractions of lysates from human coronary artery smooth muscle cells transfected with siControl or siMiro1 for 72 h. (B) Quantification of immunoblot signal in samples as shown in (A). (C) Number of human coronary artery smooth muscle cells transfected with siControl or siMiro1 at 72 h after incubation with growth media in addition to PDGF (20 ng/ml) or control. (D) Confocal images of human coronary artery smooth muscle cells grown on CYTOOchips^TM^. Cells were transfected with siControl or siMiro1 for 72 h before being seeded onto CYTOOchips^TM^. The cell cycle was synchronized by serum starvation for 24 h (not shown); the cells were subsequently treated with medium containing 10% FCS and PDGF (20 ng/ml) for 6 h. (E) Mitochondrial probability map. The cumulative distribution of mitochondria was assessed for images as in (D) by modified Sholl’s analysis. (F) Mito^95^ values, defined as the distance from the center of the CYTOOchips^TM^ at which 95% of the mitochondrial signal is found, under the conditions used in (D). (G) ATP levels assessed by luminescence assay in human coronary artery smooth muscle cells transfected with siControl or siMiro1 at 72 h. (H) Images of human coronary artery smooth muscle cells transfected with siControl or siMiro1 at 72 h after transduction with an adenovirus expressing a cytoplasmic ATP sensor with red reference protein (cyto-Ruby3-iATPSnFR1.0) for 24 h. (I) Quantification of GFP/RFP ratios, indicating cytosolic ATP levels in cells shown as in (G). (J) Quantification of activity of ETC complex 1 in human coronary artery smooth muscle cells following transfection with siControl or siMiro1 as determined by the decrease in the rate of absorbance at 340 nm with and without rotenone incubation for 10 min. (K) Quantification of activity of ETC complex 1, plotted as the difference between absorbance curve slopes with and without rotenone (as in I). Statistical analyses were performed by Mann-Whitney (B, F, H, J), Kruskal-Wallis (C) and Mann-Whitney test (G).

### Pharmacological reduction of MIRO1 impairs VSMC proliferation

Recently, a small-molecule MIRO1 reducer [20] was developed that removes MIRO1 from mitochondria. The rationale behind its development lies in the observation that MIRO1 accumulates on depolarized mitochondria in a large cohort of Parkinson’s disease fibroblasts. This reducer effectively lowered MIRO1 levels and rescued neurodegeneration in Drosophila models and in vitro studies. Notably, the genetic deletion of MIRO1 induces large, hyperfused mitochondria and delays mitophagy [29]. Though this study does not primarily focus on mitophagy as a mechanism, we evaluated whether this compound could be used to inhibit cell proliferation and ATP production. In wild type VSMCs, incubation with this compound for 72 h reduced mitochondrial mass (**Figure 8A-C**). It also decreased the number of proliferating VSMCs at 72 h **(Figure 8D**). The effect of the reducer on cell number was dose-dependent (**Figure 8E**). Furthermore, the compound also lowered intracellular ATP levels in wild type VSMCs (**Figure 8F**). These findings suggest that reducing MIRO1 could be utilized to treat vasoproliferative diseases, such as neointima formation.

**Figure 8.**
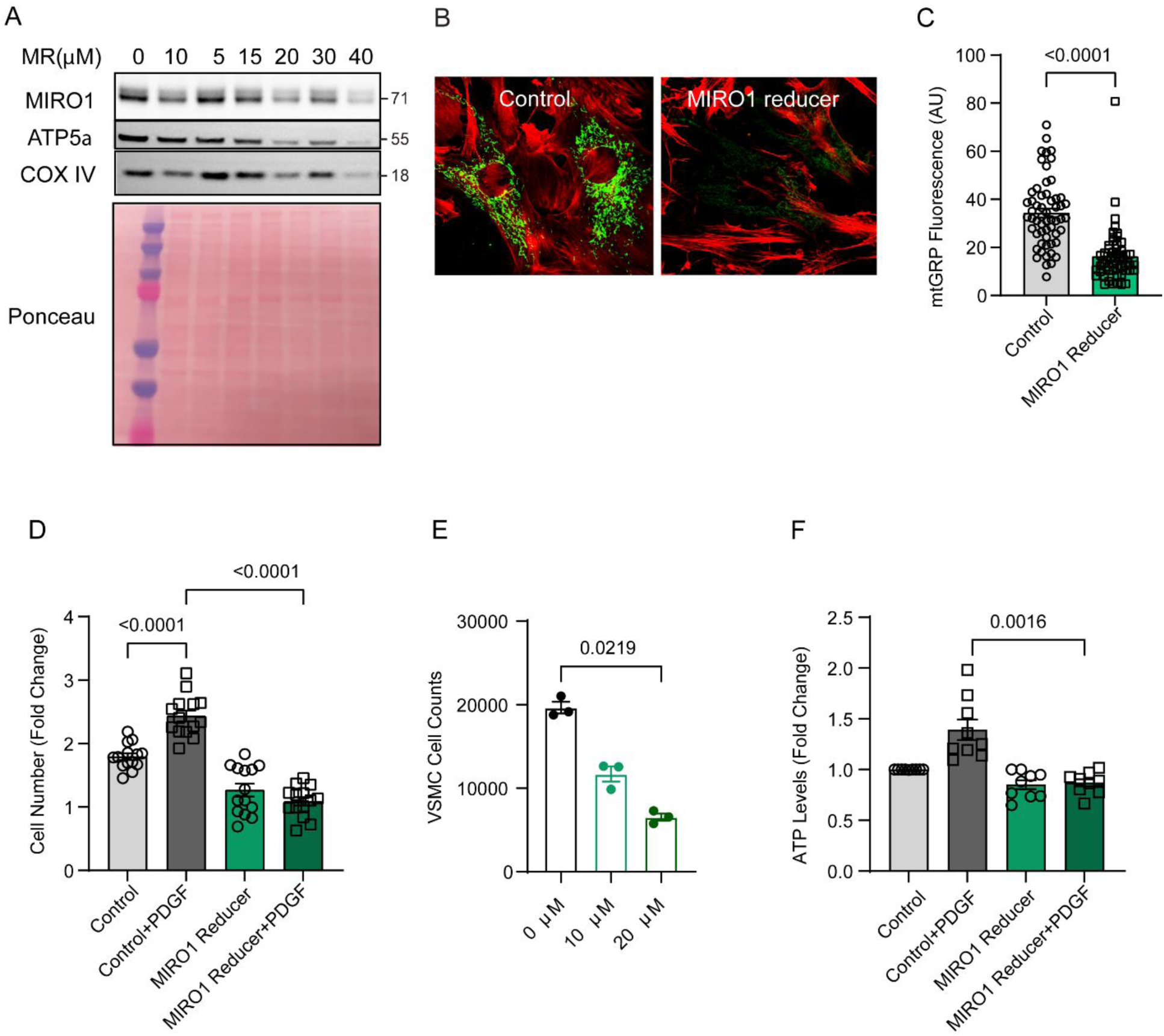
MIRO1 reducer inhibits VSMC proliferation. (A) Immunoblots for MIRO1, ATP5α and COX IV in lysates of VSMCs treated for 72 h with MIRO1 reducer (MR, concentrations as indicated). Ponceau-stained membrane as loading control. (B) Representative confocal images of wild type VSMCs treated with DMSO (control) or MIRO1 reducer (MR, 10 μM) for 48 h. (mtGFP, green; phalloidin, red). (C) Quantification of mtGFP fluorescence in experiments like those shown in (B) (n=4). (D) Cell counts of wild type VSMCs treated with DMSO or MIRO1 reducer (10 μM), following a 72-h incubation in medium containing 10% FBS with or without PDGF (20 ng/ml). (E) Cell counts of wild type VSMCs treated with DMSO or MIRO1 reducer (10 and 20 μM), (n=3). (F) ATP levels in wild type VSMCs treated with DMSO or MIRO1 reducer (10 μM), following 72-h incubation in medium containing 10% FBS with or without PDGF (20 ng/ml). Analysis by Mann-Whitney (C), one-way ANOVA (D), Kruskal-Wallis test (E) and Friedman test (F).

## DISCUSSION

This study uncovers the crucial role of the outer mitochondrial membrane GTPase, MIRO1, in driving VSMC proliferation and neointima formation. The findings underscore that mitochondrial motility, regulated by MIRO1, is essential not only for cell migration, as previously established [15, 30], but also for smooth muscle cell proliferation. Contrary to the belief that mitochondrial motility is required to align mitochondria before mitosis, the study demonstrates that proliferation is hindered in the early cell cycle stages, particularly during the ATP-demanding G1 to S phase transition. MIRO1^-/-^ cells have reduced ATP levels, emphasizing the importance of MIRO1-mediated ATP production. Based on our recent report [25], we posit that mitochondrial motility facilitates the formation of MERCS, which support enhanced calcium transfer during the G1 to S phase, ultimately boosting ATP production. Furthermore, MIRO1 forms directly associates with the MIB/MICOS complex, the microtubule motor machinery, and the ETC subunit Nduf9, suggesting a dual role in supporting both mitochondrial ATP production and motility. This insight into the interplay between mitochondrial dynamics and cellular energy metabolism refines our understanding of conserved mitochondrial mechanisms that regulate cell proliferation.

We first discovered that neointima formation is abolished in a mouse model with VSMC-specific MIRO1 deletion, and that cell proliferation in cultured aortic VSMCs lacking MIRO1 is significantly reduced due to delays in the early phases of the cell cycle. While most previous studies primarily focused on the role of MIRO1 in neurons and embryonic fibroblasts [16, 19–21, 31–35], our findings extend its importance to primary proliferating VSMCs, revealing its critical involvement in VSMC proliferation and neointima formation. We believe this is the first demonstration that MIRO1 deletion in VSMCs abolishes neointima formation in a mouse model, highlighting MIRO1 as a potential new therapeutic target for treating vasoproliferative diseases. In immortalized cells lacking MIRO1 and 2[16], impaired cytokinesis with asymmetric partitioning of mitochondria to daughter cells was reported in M-phase [36]. In addition to reduced progression in the G2/M phase in MIRO1^-/-^ VSMCs, we found robust evidence for impairment in the early phases of the cell cycle.

The loss of MIRO1 reduced intracellular ATP levels and abolished mitochondrial motility. Inhibiting mitochondrial respiration blocked their motility, whereas halting motility by microtubule disassembly did not affect ATP levels. Thus, MIRO1 not only controls motility, but also supercomplex formation and ETC activity. Our data support a model in which mitochondrial energy production is required for mitochondrial motility. Whether reduced ATP production negatively impacts mitochondrial motility because of impaired activity of the C-terminal GTPase domain of MIRO1 as recently described or impaired actin polymerization will require further investigation [37].

VSMCs are believed to mostly rely on glycolysis to fuel their proliferation [38]. However, recent studies have provided evidence that, analogous to proliferating cancer cells, VSMCs also depend on glutaminolysis [39] [40], which shuttles metabolites to the TCA cycle and promotes oxidative phosphorylation. Our findings indicate that VSMCs require mitochondrial ATP since ATP synthase inhibition blocked mitochondrial motility and VSMC proliferation. In our study, halting mitochondrial motility by microtubule disassembly did not decrease overall intracellular ATP levels in VSMCs. These data further support that mitochondrial motility controlled by MIRO1 enables subcellular mitochondrial positioning within VSMCs, which then supports localized subcellular ATP and ROS production. MIRO1 ensures that mitochondrial metabolites exert their effects on local targets, similar to findings in embryonic fibroblasts [41, 42].

We found that the reconstitution of MIRO1^-/-^ VSMCs with MIRO1-WT normalized cell proliferation, ATP levels, and motility, whereas the reconstitution of MIRO1-KK, which lacks EF hands, resulted in only partial recovery. These results suggest that the EF hands of MIRO1 are important for its full functionality in VSMCs. In neurons, MIRO1 regulates the anterograde transport of mitochondria along microtubules [16, 43, 44]. Mitochondrial arrest in response to elevated Ca^2+^ in subcellular domains and synaptic activity is driven by the Ca^2+^-sensing EF hands [32, 45]. Here, we provide evidence that without Ca^2+^-controlled mitochondrial arrest through point mutations of the EF hands, VSMC proliferation is reduced. We speculate that the loss of functional EF hands impairs alignment of mitochondria and ER. This model is supported by our recent study, in which we demonstrated that MIRO1 deficiency disrupts MERCS formation and mitochondrial calcium uptake during the cell cycle [25].

MIRO1 loss caused an intracellular energy deficit, increased AMP kinase phosphorylation, and reduced the expression of proteins promoting the G1/S cell-cycle transition. This finding suggests a mechanism where MIRO1 influences cell proliferation through its impact on cellular energy status and cell cycle regulatory proteins. Indeed, a link between AMP kinase activation and upregulation of p53 and G1/S cell cycle arrest has been described [27, 46]. Here, our data provides an explanation for how low ATP levels are caused by MIRO1 deletion and block cell cycle progression at this cell cycle stage.

We observed that loss of MIRO1 impaired the formation of mitochondrial cristae as observed in embryonic fibroblasts [21]. In our model of VSMCs with MIRO1 deletion, this correlated with reduced formation of ETC supercomplexes and activity of ETC complex I. In agreement with a prior study, MIRO1 associated with MIC19, MIC60 and SAM50 and the expression of MICOS/MIB proteins were not impaired with MIRO1 deletion [21]. In addition, we detected a previously unrecognized association between NDUFA9, a subunit of ETC complex I, and MIRO1, potentially modulating ETC complex I activity. We also detected evidence for decreased ETC complex V activity. These findings provide a potential mechanism by which MIRO1 controls mitochondrial ATP production, in addition to its putative role in decreasing metabolite transfer at distorted mitochondrial ER contact sites [16, 21, 25] .

Some findings of our study agree with published data on MIRO1, which have highlighted its role in mitochondrial dynamics, energy production, and cell signaling. Previous studies have shown that MIRO1 is involved in the regulation of mitochondrial transport and positioning, which are critical for metabolic signaling [13, 15, 41, 47]. The observed effects of MIRO1 loss on ATP production and mitochondrial structure in our study align with reports indicating that MIRO1 dysfunction can lead to impaired mitochondrial respiration and energy deficits [15, 21, 41, 44]. Additionally, the role of MIRO1 in cell proliferation is supported by the literature demonstrating its involvement in mitotic spindle formation at later stages of the cell cycle, and more recently, in the G1/S phase [15, 16, 21].

Despite these compelling findings, our study has limitations. One limitation is the use of a mouse model, which, while informative, may not fully replicate human VSMC behavior. We selected the complete ligation model for this study due to its high reproducibility and excellent surgical feasibility [48], which allows for consistent outcomes across investigators. While it does not fully replicate some aspects of human balloon angioplasty with restenosis, such as endothelial denudation, it provides valuable insights into mechanisms of neointima formation.

Additionally, our experiments focused primarily on cultured aortic VSMCs, which may not entirely represent the in vivo environment. Future studies should include a broader range of VSMC models and in vivo validations to confirm these mechanisms. Moreover, MIRO1 shares 60% sequence identity with its paralog MIRO2 [31]. Both are ubiquitously expressed in eukaryotes, yet they are not functionally redundant [16, 44]. In VSMCs with MIRO1 deletion, we did not detect a compensatory upregulation of MIRO2. The functions of MIRO2 in the vasculature are currently unknown. However, a previous study proposed that after trans-aortic banding, forced expression of MIRO2 in cardiac myocytes improved mitochondrial function [49].

In conclusion, our study demonstrated that MIRO1 is a pivotal regulator of VSMC proliferation and neointima formation. We identify specific mechanisms by which MIRO1 regulates ATP production and mitochondrial motility, linking these processes to cell cycle progression and energy homeostasis. This connection between mitochondrial dynamics and cell proliferation is a significant advancement in understanding how mitochondrial dysfunction can contribute to vascular disease. The findings suggest that targeting MIRO1 could be a potential therapeutic strategy for treating vasoproliferative diseases, such as neointima formation. Further research is needed to fully understand the downstream effects of MIRO1 loss and to explore the potential clinical applications of these insights.

## METHODS

### Data Availability

In accordance with the Transparency and Openness Promotion Guidelines, the authors declare that all supporting data and Supplemental Material are available from the corresponding authors upon reasonable request.

### Animals

Animal studies were conducted in accordance with the NIH *Guide for the Care and Use of Laboratory Animals*, following protocols that were approved by the Institutional Animal Care and Use Committee (IACUC) of the University of Iowa (#7061189 and #0051189). SMMHC-CreERᵀ² mice, in which expression of tamoxifen-inducible Cre recombinase is driven by the smooth muscle myosin heavy chain (SMMHC) promoter, were obtained from Jackson Laboratory (Strain #019079). The Mitochondrial Rho1 GTPase LoxP (Floxed Miro1 conditional KO) mice were graciously provided by Dr. Janet Shaw (University of Utah) and backcrossed with C57BL/6J mice (Jackson Laboratories, Strain #000664). The VSMC-selective floxed Miro1 conditional knockout mice (SM-Miro1^-/-^ mice) were generated by crossbreeding with SMMHC-CreERᵀ² mice. Cre recombination was induced by intraperitoneal injection of tamoxifen (80 mg/Kg) for 5 days, followed by a 14-day break. Genotyping was performed using primers 5’-CCCTGTGTCGCTGAGGTTGGAAGCTG (sense), 5’-GAAATGCCACCAGAATCCAGTGGC (sense -exon2) and 5’-GTGGAGGCAGGAGGATCAGGAG TTTAAAGTC (antisense). SMMHC-CreERᵀ² mice were injected with same dose of tamoxifen and used as a control for in vivo experiments. In vivo experiments were performed in male mice because the tamoxifen-inducible Cre recombinase in this model, whose expression is driven by the SMMHC promoter, is present on the Y-chromosome. Comparisons in this study were made between mice with and those without MIRO1 deletion in smooth muscle cells. One mouse was regarded as one experimental unit.

### Ligation of the common carotid artery and preparation of vessels for quantification of neointima and media

10–12-week-old SM-Miro1^-/-^ mice and controls were fed high-fat chow for 3 weeks (D12492, 60 kcal% fat, Research Diets) and then, the animals were anesthetized, and survival surgeries of left common carotid artery ligation were performed [48]. A high fat diet was used to approximate human-like cholesterol levels and model metabolic conditions. This diet elevates serum cholesterol to approximately 250 mg/dL and slightly increases nonfasting glucose levels (∼200 mg/dL), creating a metabolic profile close to that observed in humans with coronary artery disease (see https://www.jax.org/jax-mice-and-services/strain-data-sheet-pages/phenotype-information-380050). 21 days after the surgery, the animals were anesthetized and underwent transcardiac perfusion with 10 ml of PBS, followed by fixation with 10 ml of 4% paraformaldehyde (PFA), at physiological pressure. The left (ligated) and right (non-ligated) common carotid arteries were excised at the carotid bifurcation and embedded for sectioning followed by Verhoeff Van Gieson (VVG) staining. Images were captured using a Nikon eclipse TS100. NIH Image J was used to trace the circumference of the lumen, internal elastic lamina (IEL), and external elastic lamina (EEL), and both the area and perimeter were calculated.

Neointimal areas were calculated by subtracting the area of the lumen from that of the internal elastic lamina, and medial areas were determined by subtracting the area of the internal elastic lamina from that of the external elastic lamina. Eight mice in the control group and eight mice in the experimental group (MIRO1 deletion in smooth muscle cells) underwent ligation for a total of 16 mice. The non-ligated contralateral carotid artery of each mouse was used as control.

Treatments were performed in control and SM-Miro1^-/-^ mice at the same time to avoid potential confounders. The group size was estimated based on previous experience with this procedure. No mice were excluded from the analysis. The primary outcome was neointimal area. Two scientists who were aware of the group allocation collaboratively performed the surgeries, one processed the tissue and performed staining, imaging, and image analysis.

### Culture of vascular smooth muscle cells (VSMCs)

Primary mouse aortic VSMCs from mice on normal chow were isolated enzymatically; incubation in elastase (1 U/ml for 10 min at 37°) was performed to remove the adventitia. The medial layer of the aorta, which contains the VSMCs, was minced and incubated in 2 mg/ml collagenase type II digestion solution (Worthington Biochemical Corporation) for 2 h. Aortic VSMCs were plated in standard VSMC medium (DMEM medium containing 1% penicillin/streptomycin, MEM nonessential amino acids, MEM Vitamin and 8 mmol/L HEPES) supplemented with 20% fetal bovine serum (FBS) and 1% fungizone. The proliferative VSMCs were cultured and propagated in standard VSMC medium with 10% FBS at 37°C in humidified 95% air and 5% CO_2_ incubator. In early experiments, VSMCs were isolated from SM-Miro1^-/-^and littermate mice (referred to as SM-Miro1^-/-^ VSMCs). In later experiments, VSMCs were isolated from Miro1^fl/fl^ mice and had been transduced with adenovirus expressing Cre recombinase at a multiplicity of infection (MOI) of 50 for 2 weeks (referred to as MIRO1^-/-^VSMCs). VSMCs from littermate controls were subjected to the same procedure with empty vector control adenovirus.

### Cell lysis and fractionation

Whole cells were lysed in RIPA buffer (20 mM Tris, 150 mM NaCl, 5 mM EDTA, 5 mM EGTA, 1% Triton X-100, 0.5% deoxycholate, 0.1% SDS, pH 7.4) supplemented with both protease inhibitors (Mini complete, Roche) and phosphatase inhibitors (PhosSTOP, Roche). Lysates were sonicated and debris was pelleted by centrifugation at 10,000 × g for 10 min at 4°C. Mitochondrial fractions were prepared in MSEM buffer (5 mM MOPS, 70 mM sucrose, 2 mM EGTA, 220 mM Mannitol, pH 7.5 with protease inhibitors), with homogenization performed in cold MSEM buffer in a Potter-Elvehjem glass Teflon homogenizer (50 strokes). Nuclei and cell debris were pelleted by centrifugation at 600 × g for 5 min at 4°C. Mitochondria were separated from the cytosolic fraction by centrifugation at 8000 × g for 10 min at 4°C. Protein concentrations were determined using the Pierce™ BCA protein assay (Thermo Scientific).

### Construction and transduction of the adenoviral Miro1 cDNA

The Miro1 plasmid constructs pRK5-myc-Miro1, pRK5-myc-Miro1 E208K/E328K and pRK5-myc-Miro1 Δ593 – 618 were obtained from Addgene (47888, 47894, 47895). The full-length ORF of the Miro1 gene (GenBank ID: AJ517412) was subcloned into the adenoviral shuttle pacAd5-CMV-mcs SV40pA (University of Iowa Viral Vector Core) by incorporation of 5’ ClaI and 3’ EcoRI restriction sites to facilitate ligation specifically. Positive clones were identified by restriction digestion. The viral constructs, E208K/E328K mutants, and N-terminal Myc tag were verified by DNA sequencing with the forward primer 5’-GTG GGA GGT CTA TAT AAG CAG AGC TCG-3’ and the reverse primer 5’-TTA AAA AAC CTC CCA CAC CTC CCC-C-3’. Additionally, expression of the MIRO1 protein and the integrity of Ad5 shuttle were confirmed before adenoviral particles were packaged, and virus was titrated using the RapAd™ system developed by the University of Iowa Viral Vector Core Facility. An MOI of 100 was determined to be sufficient to efficiently transduce the mouse VSMCs in serum-free medium overnight. The same strategy was used to subclone the fluorescent ATP sensitive protein pm-iATPSnFR1.0 subcloned into the adenoviral shuttle pacAd5-CMV-mcs SV40pA to generate adenoviral particles.

### Miro1 knockdown in human coronary artery smooth muscle cells

Human coronary artery endothelial cells (HCAECs) were kindly provided by Dr. Gerene Denning and Dr. Neal L. Weintraub. Cells were seeded in T75 cm cell culture flasks in DMEM medium, which was supplemented with 10% FBS, 1% penicillin/streptomycin, MEM non-essential amino acids, MEM Vitamin Solution, and 8 mmol/L HEPES at 37°C in a humidified, 95% air/5% CO_2_ incubator. To assess MIRO1 knockdown, siRNA duplexes 5’-GUUGUUGCAGAUAUCUCAGAAUCG-3’ and 5’- UCAGUGUCAGCUUACCAACAUGACA-3’ (IDT hs.RiRHOT1.13.1 and 3) or a scrambled negative control (5’-CUUCCUCUCUUUCUCUCCCUUGUGA-3’) were used. Transfection was performed using Lipofectamine RNAiMAX (Invitrogen Cat# 56532) based on the manufacturer’s protocol. Briefly, 20 nM of scrambled or MIRO1 siRNAs and 10 µL of lipofectamine RNAiMAX were incubated together in 500 µL of Gibco Opti-MEM (Thermofisher Scientific, Cat# 31985-070) at room temperature for 30 min. Transfection was performed in a low volume of Opti-MEM for 5 hours before supplementation with DMEM media. Knockdown efficiency was evaluated by immunoblotting 72 hours post-transfection.

### Cell counting

For calculations of the average proliferation rate of a population, the wells of 12-well plates were seeded with 10,000 cells or 35mm dishes with 40,000 to 70,000 cells, and cells were counted at multiple cell confluency points. After seeding VSMCs for 24 h, they were treated with PDGF (20 ng/ml) and cultured for additional 72 h. Cells were dissociated with Gibco™ TrypLE Express and counted in triplicate using a Beckman Coulter Z1 cell counter. In some experiments, Oligomycin (10 µM) or MIRO1 reducer (10 and 20 µM) were added to PDGF.

### Flow cytometry analysis of cell cycle progression

VSMCs were initially synchronized in G0/G1 phase by serum starvation for 48 h and this arrest was terminated by adding 10% FBS medium for 0, 24 and 48 h, respectively. 1 x 10⁷ cells were harvested and fixed with cold 80% ethanol for 1 h, treated with 1 mg/ml DNase-free RNase A (Thermo Scientific), and stained with propidium iodide solution (100 ug/ml) overnight. Cytometry was performed using the Becton Dickinson LSR II instrument (561 nm laser). Forward scatter (FS) and side scatter (SS) were measured to identify single cells, and pulse processing (PI area vs. PI width) was used to exclude cell doublets from the analysis. The cell cycle data were acquired using the BD FACS Diva software.

### CYTOOchip™ micropattern

Cells were transduced with mito-targeted GFP (Ad5-CMV-Cox8-mtGFP), and Y-shaped micropatterned CYTOO™ chips were coated with collagen Type I (40 ug/ml). 150,000-200,000 cells were plated per chip, according to the manufacturer’s protocol. The cells were allowed to attach to and spread on the adhesive micropatterned substrate for 8 h in 10% FBS VSMC growth medium, at which point the medium was replaced with the serum-free VSMC growth medium for 16 h. After the synchronization protocol was completed, the cells were treated with 10% FBS/PDGF-BB (10 ng/ml) for 6 h, fixed with 4% PFA, stained with Alexa Fluor 568 Phalloidin to label F-Actin, and imaged. Images were captured using a Zeiss LSM 880 confocal microscope with objective of at 63X. The mitochondria were visualized with adenoviral mtGFP (488 nm), cell shapes were visualized with phalloidin (568 nm), and nuclei were visualized with To-Pro3 (633 nm). The mitochondrial distribution was analyzed using a custom-made ImageJ plugin.

### Analysis of mitochondrial distribution based on CYTOO images

The distribution of mitochondria was analyzed with ImageJ plus the custom-made plugin written by Guillermo Lopez-Domenech in the laboratory of Dr. Joseph Kittler at University College London. An image of F-actin staining, which defines the cell structure, was used to identify the cell center. A threshold value (pixels) for the mitochondrial area was adjusted manually. The intensity of mitochondrial signal within shells radiating out from the cell center at 1 um intervals was quantified. The cumulative distribution of mitochondrial signal was normalized per cell. An average value was calculated for each distance interval within each cell. For each experimental group, a mitochondrial probability map (MPM) was plotted. The distance at which 95% of the total mitochondrial mass (Mito^95^) is found was calculated for each cell by interpolation. One average Mito^95^ value was calculated per group.

### Mitochondrial motility imaging

VSMCs were transiently transfected with mito-ds-red and untargeted GFP adenoviruses to fluorescently label the mitochondria and cytosol, respectively. Live-cell fluorescence confocal microscopy videos were acquired using a Zeiss LSM 980 confocal microscope under controlled temperature, CO_2_, and humidity levels. Videos were generated from images taken at one-minute intervals over a 25-30-minute period.

### Mitochondrial shape imaging and analysis

Mitochondrial shape was assessed in VSMCs transduced with mito-GFP adenovirus, imaged using a Zeiss LSM 510 confocal microscope (magnification 40X). Mitochondrial form factor, defined as the ratio of mitochondrial length to width, was quantified using a custom-written NIH ImageJ macro provided by Dr. Stefan Strack (University of Iowa) [50, 51].

COX1 forward TGCTAGCCGCAGGCATTAC, reverse GGGTGCCCAAAGAATCAGAAC mtRNR1 forward GGGATTAGATACCCCACTATGCTT, reverse CCGCCAATGCCTTTGAGTTTTAAG NDUFV1 forward CTTCCCCACTGGCCTCAAG, reverse CCAAAACCCAGTGATCCAGC

### Transmission electron microscopy and analysis of the mitochondrial ultrastructure

VSMCs were washed twice with 1x Dulbecco’s phosphate-buffered saline (DPBS) and fixed overnight at 4°C, in 2.5% glutaraldehyde prepared in 0.1M sodium cacodylate buffer [pH 7.4; EM grade]. Following primary fixation, samples were washed three times in the same buffer and post-fixed with 1 % osmium tetroxide for 1 hour at room temperature. The samples were dehydrated by submersion in a series of ethanol concentrations (50%, 70%, 90%, 95%, to 100%) and then infiltrated with Epon 812 resin (Ted Pella, Redding, CA). Polymerization was conducted at 70°C overnight. Ultrathin sections (∼50 nm) were obtained using a Leica EM UC7 ultramicrotome and collected on formvar/carbon-coated copper grids. Sections were subsequently stained with 2.5% uranyl acetate and lead citrate for 2 min. Imaging was performed using a JEOL JEM-1230 transmission electron microscope (JEOL, Peabody, MA) and photographed using a Gatan UltraScan 1000 2k x 2k Tietz CCD camera (Gatan, Pleasanton, CA) at the Central Microscopy Research Facility at the University of Iowa.

### Systematic Quantification of Morphology

Using previously documented parameters and quantification methods [52–55], a unique and trained experimenter imaged the entire cell at low magnification. The obtained images were imported into ImageJ in an acceptable format, such as TIFF. The cell was then divided into quadrants using the ImageJ Quadrant Picking plugin to ensure random and unbiased quadrant selection for quantification. After dividing the images into four quadrants, two quadrants were randomly selected for complete quantitative analysis.

To minimize subjective bias, three independent, blinded quantified analysts quantified these selected quadrants as described below. Their collective findings were averaged to decrease individual subjective bias. Measurements were repeated on a minimum of 10 cells to ensure accurate and reproducible values. In the future, if significant variability is observed among the individuals performing the analysis, increasing the sample number (*n*) by expanding the number of cells quantified was found to decrease variability.

All analysis methods were developed using NIH ImageJ software. Before analysis, necessary measures should be set on ImageJ (Analyze > Set Measurements: area, mean gray value, min & max gray value, shape descriptors, integrated density, perimeter, fit ellipse, Feret’s diameter). Mitochondria were measured, including area, circularity, and length, using the multi-measure region of interest (ROI) tool in ImageJ-based on established measurements [52–55]. Then also cristae score and cristae volume. Using the freehand tool in NIH ImageJ 1.49, the mitochondria and cristae membranes were manually traced to determine the area or volume.

### Immunoblotting

Frozen tissues were homogenized using a pestle and mortar filled with liquid nitrogen. The resulting tissue powder was resuspended in four volumes of RIPA buffer containing protease inhibitor cocktail. Cells were lysed in RIPA buffer. Homogenates were centrifuged at 12,000 rpm for 20 min at 4°C. Protein in the supernatants was quantified using the Pierce BCA protein assay. Equal amounts of denatured protein were electrophoresed on Novex 4-12% SDS NuPAGE pre-cast gels (Life Technology) and transferred to Immobilon-P 0.45 μm PVDF membrane. Membranes were incubated with primary antibody in 1%BSA in TBS-T buffer at 4°C overnight, and in 1%BSA in TBS-T buffer containing the appropriate HRP-conjugated secondary antibody at room temperature for 2 h. The blots were washed thoroughly and visualized with Clarity™ Western ECL substrate, using the ChemiDoc™ Touch imaging system according to the manufacturer’s instructions, and quantitated using the Image Lab 6.0 software (Bio-Rad). The densitometry was performed using ImageJ software.

### Pull-down assays

HEK cells were infected with adenovirus encoding c-myc tagged MIRO1-WT or MIRO1-KK for 24h. c-myc mouse antibody (0.5 ug/reaction, Santa Cruz, sc-40) was precipitated using magnetic Dynabeads Protein G (25 μm /reaction, Thermofisher Scientific 10003D) following a 30 min incubation at room temperature. Specifically, the beads were washed 3 times with TBS buffer and incubated with protein lysates (1 mg) overnight at 4°C. After 3 washes, protein was eluted from magnetic beads by boiling in lithium dodecyl sulfate (LDS) for 10 min at 50°C. Samples were then run on Bis-Tris acrylic gels for immunoblotting.

### Blue-Native PAGE

Mitochondrial fractions purified from cultured VSMCs, or aortic tissue were resuspended in 50 μl of 6-aminocaproic acid (ACA) buffer containing 750 mM ACA (Sigma A2504), 50 mM Bis-Tris pH 7.0 (Sigma B9754), and 1x protease inhibitors. Protein concentrations of the samples were measured by Bicinchoninic Acid (BCA) assay, using 4 µl of the resuspended mitochondrial proteins. For BN-PAGE, 40 ug of mitochondrial protein samples were solubilized in 1.5% digitonin (Invitrogen, Cat. No. BN2006), with a final ratio of 8:1 of digitonin to mitochondrial proteins and incubated on ice for 20 min with gentle vortexing at 1000 rpm. The solubilized samples were centrifuged at 16,000 *g* for 30 min at 4°C to collect clarified proteins. The loading samples were prepared in 0.2% of NativePAGE G-250 Sample Additive (Invitrogen, Cat. No. BN2004) and glycerol. The native protein samples were resolved by NativePAGE (3-12% Bis-Tris Gel Invitrogen, Cat. No. BN1001BOX) according to the manufacturer’s protocol. Briefly, the gels were run at 4°C in chilled Dark Blue Cathode Buffer (Invitrogen, Cat. No. BN2001) at 130V for 40 min, and then in Light Blue Cathode Buffer (Invitrogen, Cat. No. BN2002) at 250V for 60 min. The NativeMark^TM^ Unstained Protein Standard (Invitrogen, Cat. No. LC0725) was used to estimate the molecular weight. The gels were rinsed with transfer buffer (190 mmol/L glycine, 27 mmol/L Tris) and the proteins were transferred to 0.45 μm PVDF membranes at 25 mA overnight in a mini transfer tank (Bio-Rad). The membranes were incubated in 8% acetic acid for 15 min and dried for 2 h. They were then reconstituted in 100% methanol for 2 min and rinsed with distilled water for 5 min before being blocked and incubated with antibody. Complex V activity assay was performed on mitochondrial fractions as described elsewhere [56].

### Antibody table

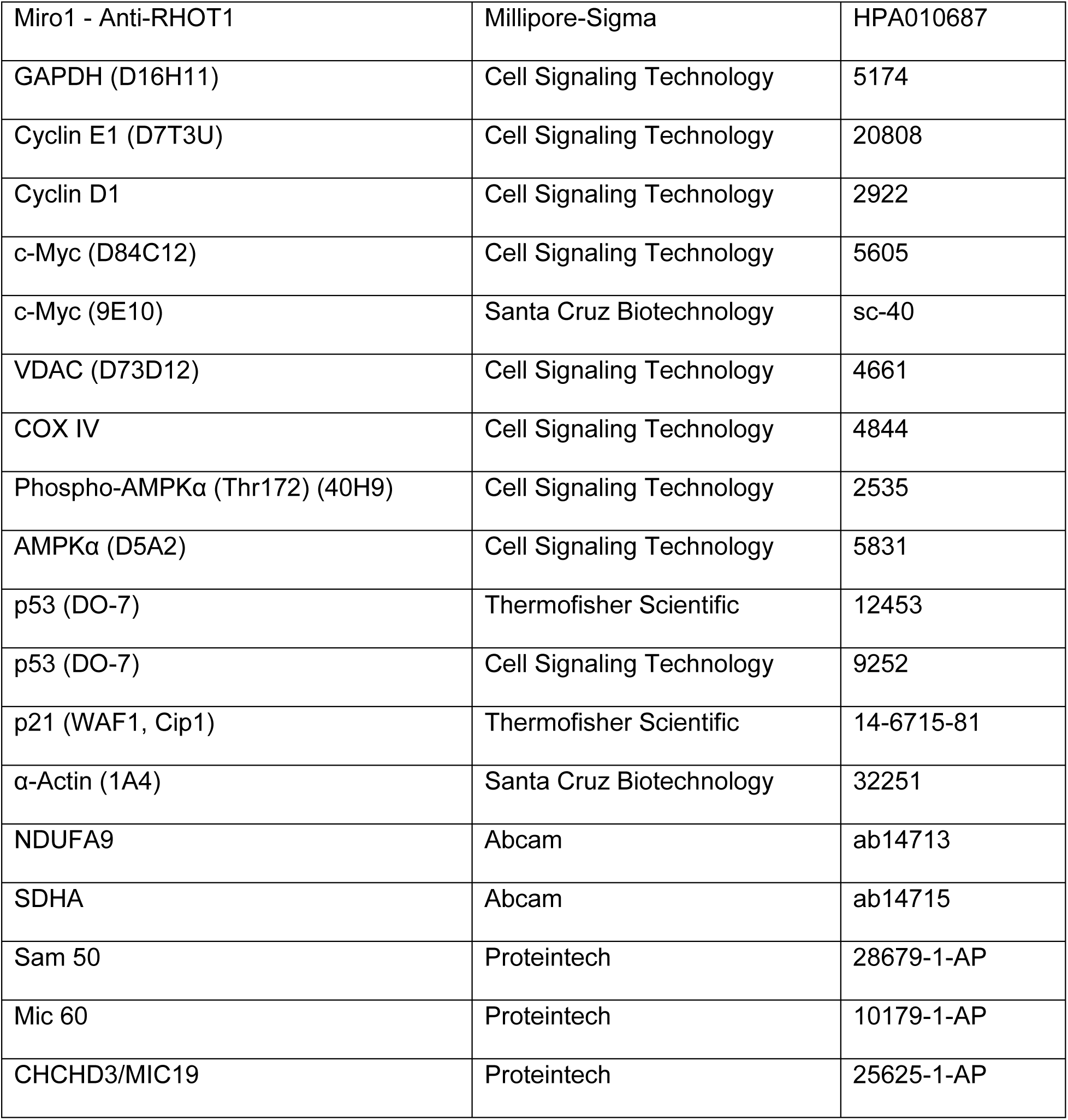
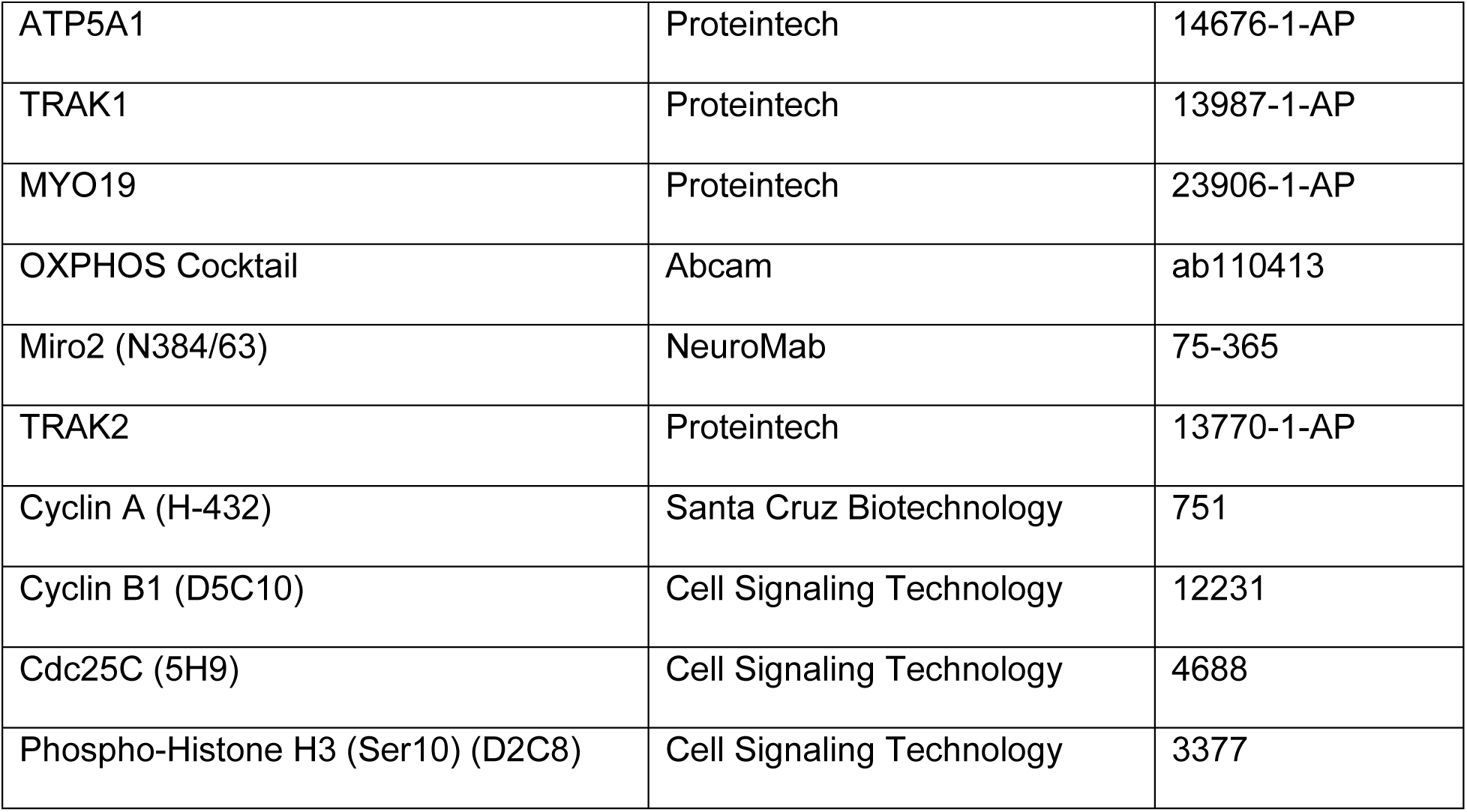

### Collection of human coronary arteries

Deidentified samples of human coronary arteries were obtained from the Iowa Living Human Heart Research Program (ILHHRP) after approval by the IRB of the University of Iowa (IRB # 200701730). The presence or absence of coronary artery disease (CAD) was determined by review of the electronic medical record system and was based on coronary angiography. All patients with CAD also had ischemic cardiomyopathy (ICM) and underwent heart transplant. Coronary arteries from explanted hearts with ICM were dissected and sections with plaques were visually identified grossly, then confirmed by microscopy. The healthy coronary artery was obtained from a patient with non-ischemic cardiomyopathy undergoing left ventricular assist device (LVAD) placement and was recovered from the portion of the LV apex removed for LVAD placement.

### Immunohistochemistry

Aortic tissues were deparaffinized, permeabilized with 0.1% Triton X-100 for 10 min at room temperature and blocked using the MOM® detection kit (Vector Laboratories). The sections were incubated for 60 min with either Anti-MIRO1 primary antibody (Cat# HPA010687, Atlas Antibodies) or anti-PCNA primary antibody, and subsequently for 30 min with an HRP-conjugated secondary antibody. DAPI was then applied for 10 min, after which the sections were counterstained with Harris hematoxylin for 10 sec. Approval for the use of coronary artery tissue from human subjects was obtained from the Institutional Review Board (IRB) of the University of Iowa.

MIRO1 expression: Human coronary artery tissue sections were blocked with 10% donkey normal serum (Jackson Immuno Research Lab., USA) for 1 h, and then incubated with rabbit anti-MIRO1 antibodies (1:10 dilution, Sigma) in 10% donkey normal serum at 25°C overnight. After they were thoroughly rinsed with PBS (phosphate buffered saline), they were incubated with affinity purified Alexa Fluor 488-conjugated anti-rabbit secondary antibody (1:200 dilution, Jackson ImmunoResearch Lab) at 4°C overnight. The sections were then rinsed with PBS and stained with a fluorescent nucleus dye (TOPRO-3,1: 2000 dilution, Molecular Probes) and a fluorescent actin dye (Alexa Fluor 568-Phalloidin, 1:40 dilution, Molecular Probes) for 15 min. The sections were rinsed and cover-slipped with Prolong Diamond Antifade Reagents (Invitrogen-Molecular Probes, USA). The stained sections were examined with a Nikon AX confocal laser-scanning microscope. Digital confocal images were obtained and processed with software provided with the Nikon AX.

MIRO1 and Ki67 colocalization: Tissue sections were incubated with rabbit anti-Miro1 antibodies (1:10 dilution, Sigma) and rat anti-Ki67 antibody (1:400, Invitrogen) in 10% donkey normal serum at 25°C overnight. They were then incubated with affinity purified Alexa Fluor 488-conjugated anti-rabbit secondary antibody and affinity purified Alexa Fluor 647-conjugated anti-rat secondary antibody (both at 1:200 dilution, Jackson ImmunoResearch Lab) at 4°C overnight. The sections were then stained with a fluorescent actin dye (Alexa Fluor 568-Phalloidin, 1:40 dilution, Molecular Probes) for 15 min.

### ATP bioluminescence assay and mitochondrial bioenergetics during G1/S phase

For assays of total ATP levels, 35mm dishes were seeded with wild type or MIRO1^-/-^ VSMCs at 50,000 cells per well, and the cells were grown in 10% FBS standard VSMC medium for 24 h. Cells were synchronized in G0/G1 phase by serum starvation for 48 h, and then the medium was replaced with 10% FBS medium/PDGF for 16 h. Cells were treated with 10% FBS medium containing 0.2% trichloroacetic acid, which protects against ATP degradation by inactivating ATPases. VSMCs were trypsinized, counted, and harvested as cell pellets that were suspended in 100 μl of 80% methanol/5mM EGTA solution for ATP determination assay. Total ATP was quantitated using the luciferin-luciferase bioluminescence ATP Determination Kit (Molecular Probes™ A22066) according to the manufacturer’s instructions. All measurements were performed in triplicate. The total ATP content in each sample was normalized based on the number of cells per well. ATP/ADP and ATP/AMP ratios were assessed using an ATP/ADP/AMP Assay Kit (Cat#: A-125) purchased from Biomedical Research Service, University at Buffalo, New York).

The oxygen consumption rate (OCR) and the extracellular acidification rate (ECAR) of live cells were determined using an Agilent Seahorse XF24 analyzer in the University of Iowa Metabolic Phenotyping Core Facility. Wells of a Seahorse 24-well XF microplate were seeded at the optimal density of 20,000 per well. Cells were grown for 24 h and equilibrated in bicarbonate-free XF calibrant medium for 1 h. PDGF (20 ng/ml) was added 16 h prior to the start of the XF assay. The basal OCR was measured before sequential addition of oligomycin (1 μl), FCCP (1.5 μl), and antimycin A/rotenone (1 μl) to induce mitochondrial stress during the assay.

### Measurement of ATP

Adenovirus from fluorescent ATP sensor plasmid (cyto-Ruby3-iATPSnFR1.0, Addgene #102551) was generated by the University of Iowa Viral Core. The plasmid expresses a cytoplasmic ATP sensor in mammalian cells with red reference protein. Wild type and MIRO1^-/-^VSMCs were infected with virus (MOI 50) 24 h before fluorescence was measured on a Nikon Eclipse Ti2 inverted light microscope. Cells were excited at 486 and 558 nm. Intensity of the GFP signal was measured at 510 nm and that of the mRuby signal was measured at 605nm. The GFP:mRuby ratio was used as an indicator of ATP levels.

### Assay of activity of Electron Transport Chain (ETC) Complex I

Mitochondria isolated from VSMCs were processed for the mitochondrial ETC Complex 1 activity Assay Kit (Cayman, 700930) using the manufacturer’s protocol. The rate of NADH oxidation was determined by monitoring the decrease in absorbance at 340 nm in the presence and absence of 1 µM rotenone, and the values were normalized to protein concentration measured by the bicinchoninic acid (BCA) assay. Differences in slopes of curves recorded with and without rotenone represented the activity of Complex1.

### Statistical Analysis

Statistical analysis was performed under guidance from the University of Iowa Department of Biostatistics. Data were analyzed using the GraphPad Prism 10.0 software and expressed as mean ± SEM. Normality and equal variance were assessed. Statistical comparisons between two groups were conducted using the unpaired t test when a normal distribution could be assumed. Otherwise, the Mann-Whiney U test was used. One-way analysis of variance (ANOVA) followed by Tukey’s multiple comparisons was used for multiple group comparisons when a normal distribution could be assumed. Otherwise, the Kruskall-Wallis test with Dunn’s multiple comparisons was used. Two-way ANOVA followed by Šidák multiple comparisons was used for grouped data sets. A *P*-value of <0.05 was considered significant, and p-values are indicated in the figures.

## Acknowledgements

The authors thank Dr. Christine Blaumueller of the Scientific Editing and Research Communication Core at the University of Iowa Carver College of Medicine for critical reading of the manuscript.

---

**Figure 1- Figure supplement 1.**
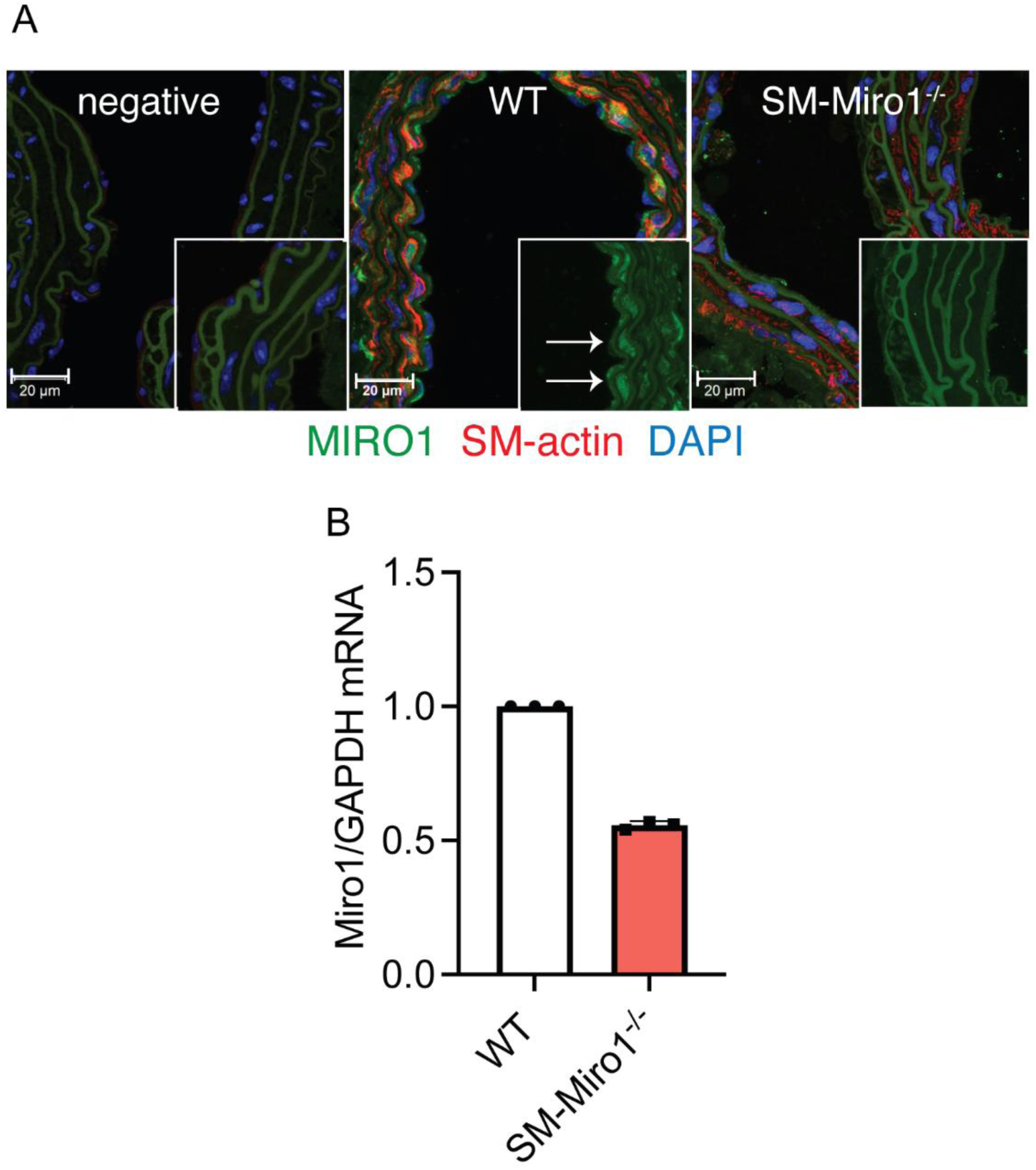
MIRO1 expression in a transgenic model of Miro1 deletion in VSMCs (SM-Miro1^-/-^). (A) Representative images of MIRO1 expression in sections of the mouse carotid artery, as determined by immunofluorescence. MIRO1 green, SM-actin red, DAPI blue. Scale bar = 20 µm. (B) Levels of the Miro1 mRNA in the murine SM-Miro1^-/-^ aorta vs wild type counterparts, as determined by quantitative real-time PCR.

**Figure 1- Figure supplement 2.**
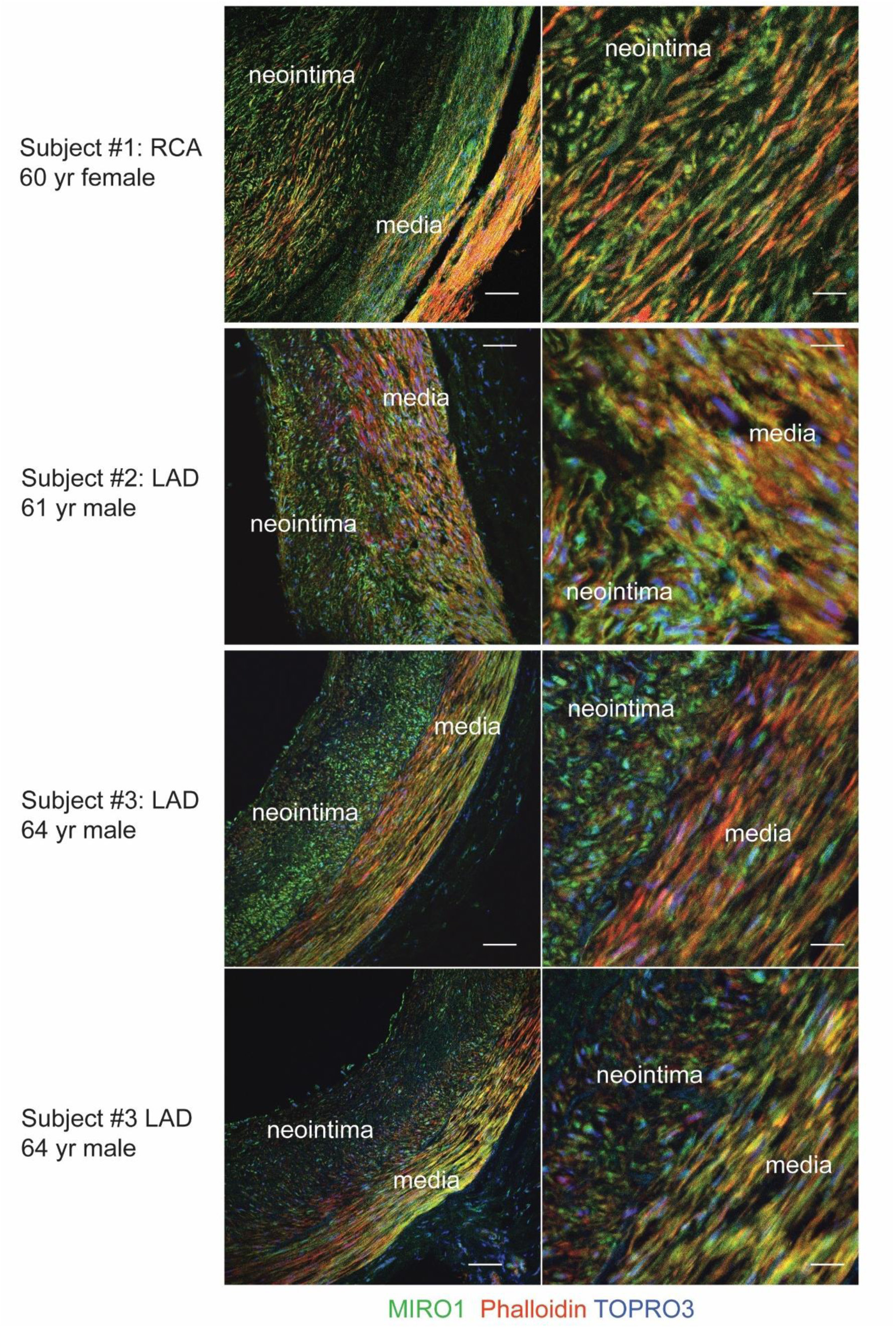
MIRO1 expression in smooth muscle cells in the human neointima. Immunofluorescence of MIRO1 in coronary arteries (left anterior descending artery, LAD; right coronary artery, RCA) of heart transplant recipients with a diagnosis of coronary artery disease. MIRO1, green; phalloidin, red; TOPRO3, blue. Scale bar: 80 µm in left images; 20 µm in right images.

**Figure 1- Figure supplement 3.**
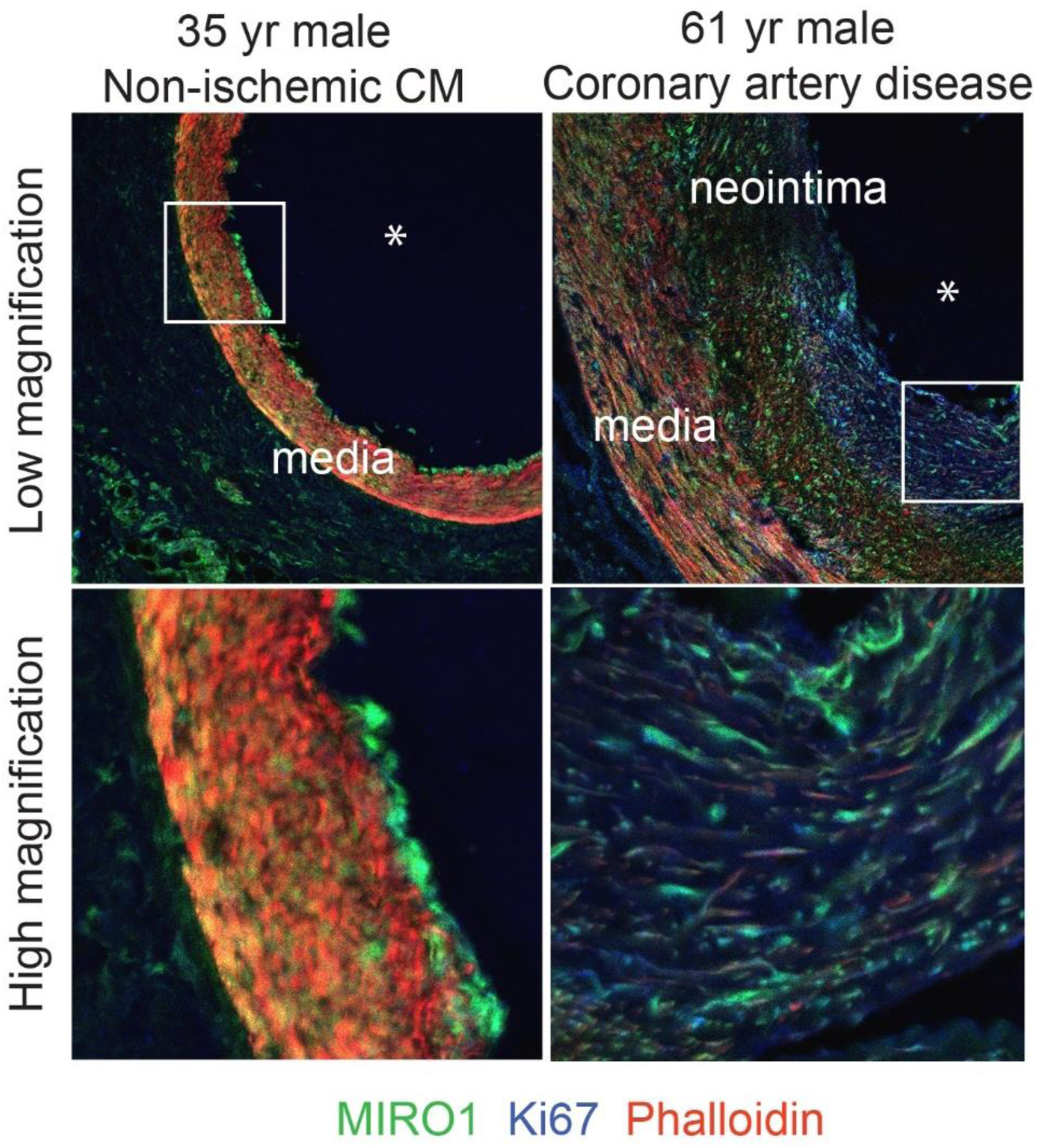
Ki67 colocalizes with MIRO1 in the human neointima. Immunofluorescence of MIRO1 and Ki67 in coronary arteries of heart transplant recipients with non-ischemic cardiomyopathy (left) and coronary artery disease (right). MIRO1, green; phalloidin, red; Ki67, blue. Same magnifications as in Figure 1- Figure supplement 2 were used.

**Figure 2- Figure supplement 1.**
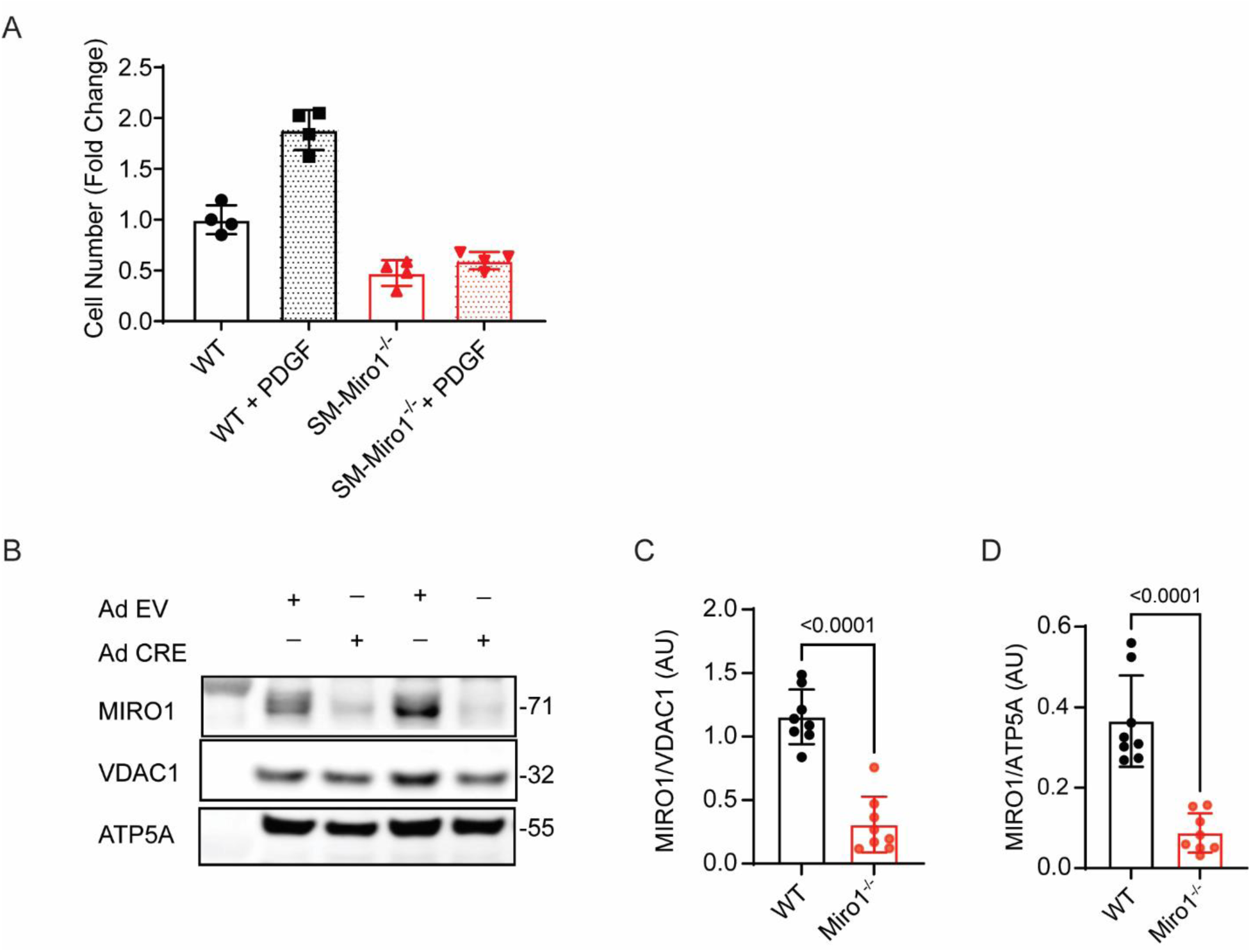
Deletion of MIRO1 inhibits VSMC proliferation. (A) Cell counts of aortic VSMCs explanted from wild type and SM-Miro1^-/-^ mice, following a 72-h incubation in medium containing 10% FBS with or without PDGF (20 ng/ml). (B) Immunoblot of MIRO1 expression in mitochondrial fractions of Miro1^fl/fl^ VSMCs transduced with adenovirus expressing cre recombinase (Ad CRE, denoted as Miro1^-/-^VSMCs) or empty vector control adenovirus (AD EV, denoted as WT VSMCs). ATP5A and VDAC1 serve as loading controls. (**C, D**) Quantification of experiments shown in (B). Analysis by unpaired t-test (C, D).

**Figure 3- Figure supplement 1.**
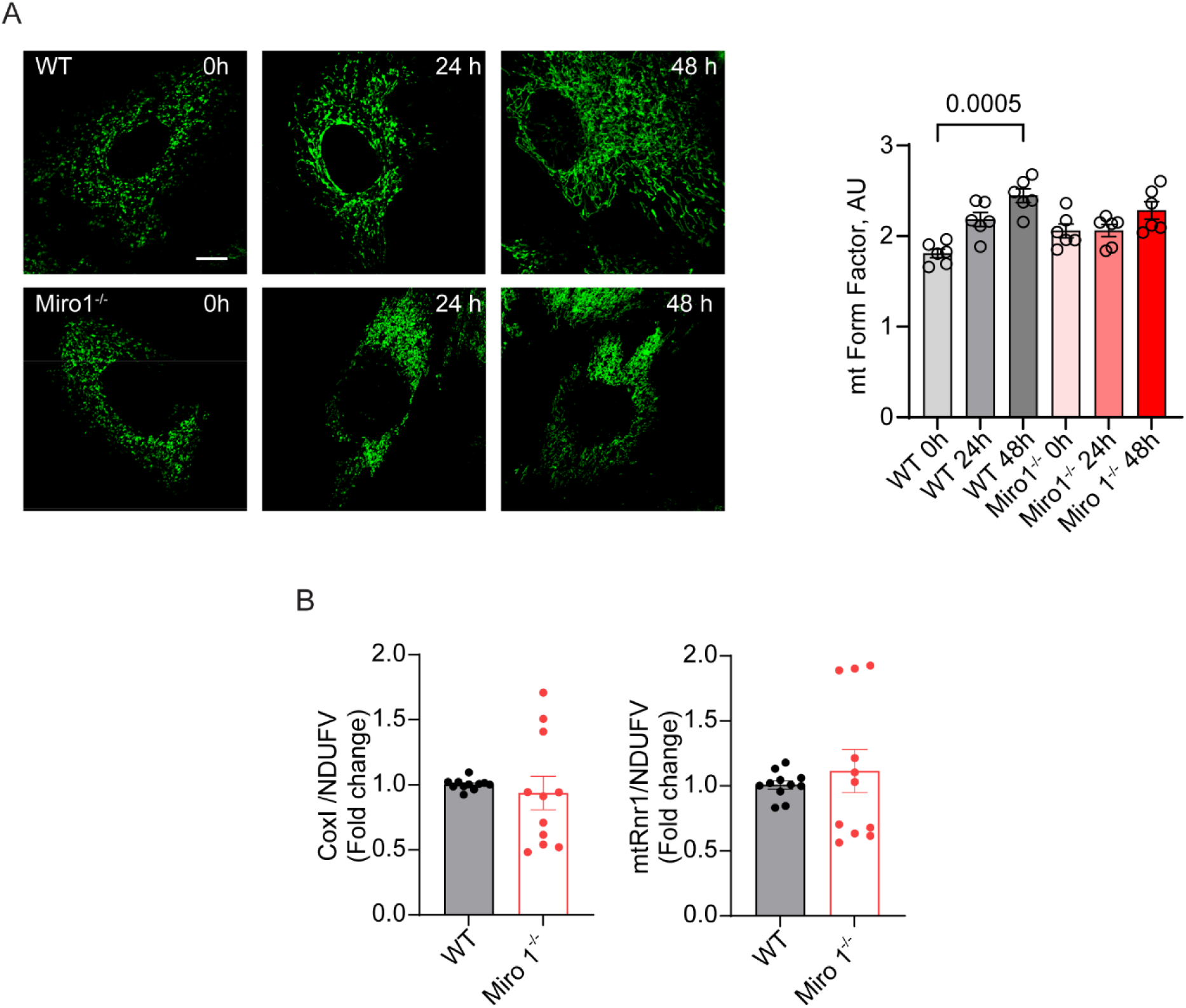
Deletion of MIRO1 alters mitochondrial dynamics without altering mitochondrial DNA copy number. **(A)** Confocal microscopy images of WT and MIRO1^-/-^ VSMCs and quantification of mitochondrial morphology as a form factor at 0, 24, and 48 h after release from growth arrest for 48 h in FBS-free media. **(B)** Quantitative polymerase chain reaction for genes encoded by mitochondrial DNA normalized to NDUFV. Statistical analyses were performed by Kruskal-Wallis (A), and Mann-Whitney (B).

**Figure 5- Figure supplement 1.**
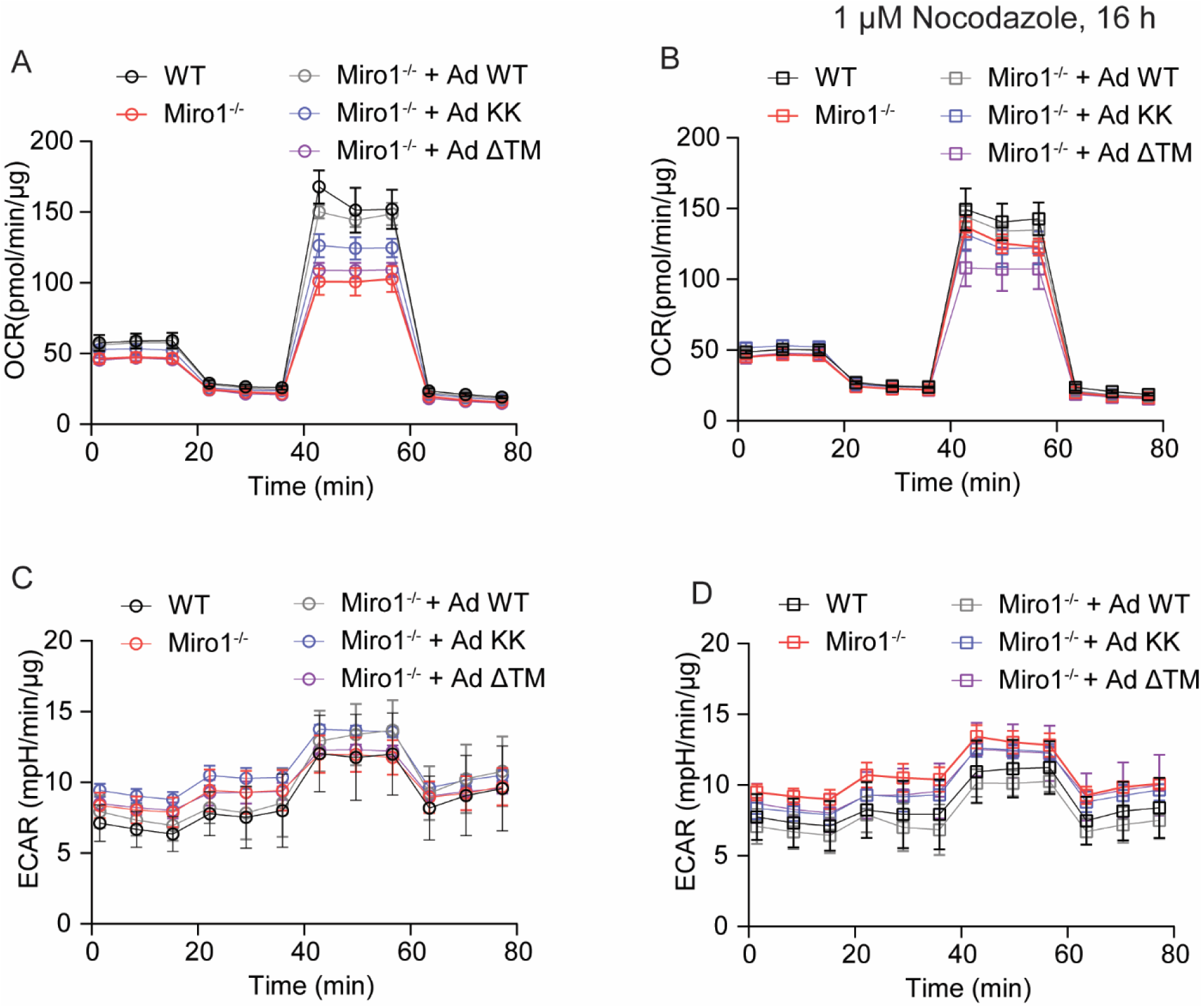
Nocodazole treatment does not impair cellular bioenergetics. (**A, B**) Oxygen consumption rate (OCR), as determined by Seahorse, for WT, Miro1^-/-^, and Miro1^-/-^ VSMCs transduced with adenovirus expressing MIRO1-WT (Ad WT), MIRO1-KK (Ad KK), or MIRO1-ΔTM (Ad ΔTM) VSMCs without (A) and with (B) Nocodazole treatment (1 µM for 16 h). (**C, D**) Extracellular acidification rate (ECAR), as determined by Seahorse, for WT, Miro1^-/-^, and Miro1^-/-^ VSMCs transduced with AdWT, AdKK, or Ad ΔTM without (C) and with (D) Nocodazole treatment (1 µM for 16 h).

**Figure 6- Figure supplement 1.**
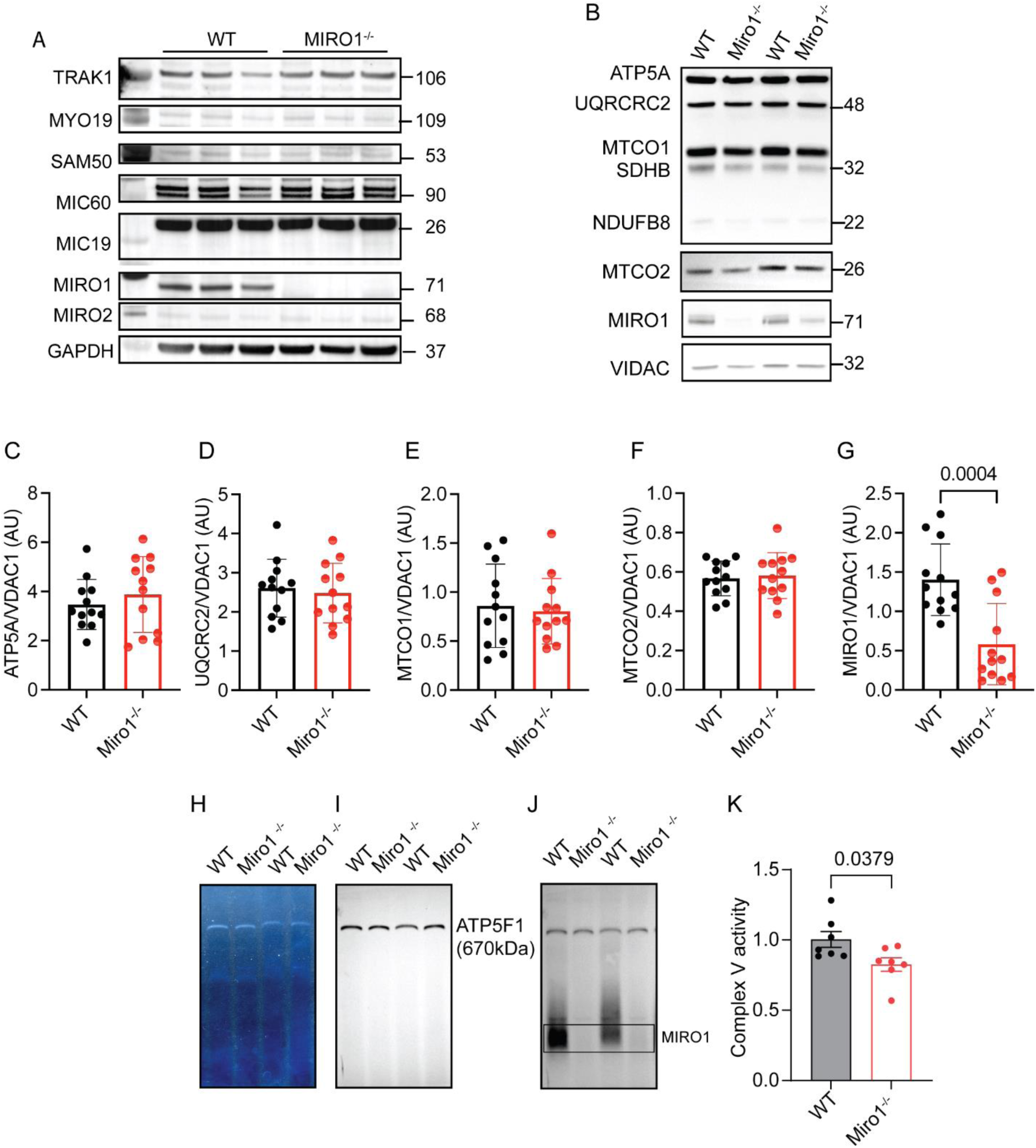
Miro1 deletion does not affect levels of subunits of MIB/MICOS or ETC complexes. (**A**) Immunoblot of MIB/MICOS proteins in whole cell lysates of wild type and Miro1^-/-^ VSMCs. (**B**) Immunoblot of subunits of ETC complexes in mitochondrial fractions of wild type and Miro1^-/-^VSMCs. (**C-G**) Quantification of (C) ATP5A, (D) UQCRC2, (E) MTCO1, (F) MTCO2, and (G) MIRO1 normalized to VDAC1 levels. (n=12 independently isolated samples per genotype). **(H-J)** Complex V in gel-activity assay in mitochondrial samples from wild type and Miro1^-/-^VSMC (H), normalized to ATP5F1 expression levels (I). Miro1^-/-^ knockout validation (J). **(K)** Quantification of Complex V activity as in (H). Statistical analyses were performed by unpaired t-test (C-G, K).

**Figure 6- Figure supplement 2.**
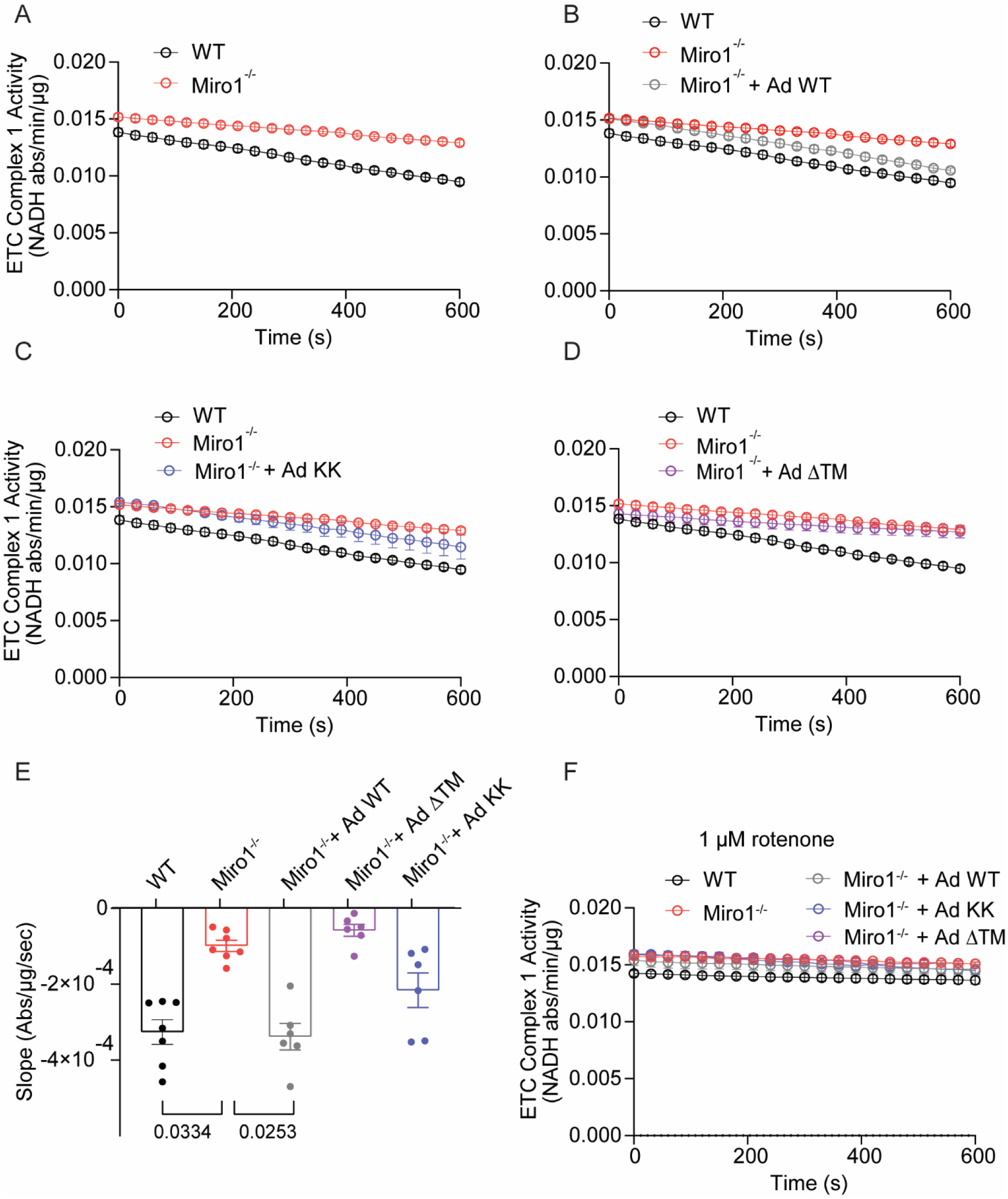
Complex I activity is regulated by the wild type form of MIRO1. (**A-D**) Quantification of activity of ETC complex 1 in WT and Miro1^-/-^ VSMCs (A), WT, Miro1^-/-^, and Miro1^-/-^ VSMCs transduced with adenovirus expressing with MIRO1-WT (Ad WT) (B), MIRO1-KK (Ad KK) (C), or MIRO1-ΔTM (Ad ΔTM) (D) as determined by the decrease in the rate of absorbance at 340 nm. (**E**) Quantification of activity of ETC complex 1, plotted as the difference between absorbance curve slopes (as in A-D). (**F**) Quantification of activity of ETC complex 1 in WT and Miro1^-/-^ VSMCs, WT, Miro1^-/-^, and Miro1^-/-^ VSMCs infected with Ad WT, Ad KK or Ad ΔTM with rotenone. Statistical analyses were performed by Kruskal-Wallis (E).

